# A bivalent lysine-acetylated small-molecule binding site in MYC

**DOI:** 10.64898/2026.03.02.707789

**Authors:** Dikshat G. Gupta, Mihai I. Truica, Jin-gon Shim, J. Brandon Parker, Mahil Lambert, Adam W. T. Steffeck, Daniel H. Ryan, Hao Pan, Kerem Kahraman, Xinyan Lu, William Yang, Kenji Unno, Ala A. Elmashae, Megan M. Kerber, C. Elysse Brookins, Huiying Han, Mary Fiona Dufficy, Songhua Quan, Debabrata Chakravarti, Gary E. Schiltz, Joshua J. Ziarek, Sarki A. Abdulkadir

## Abstract

MYC is an important, yet challenging target in oncology as it lacks traditional druggable pockets. Here, we show that two regions of MYC, the basic-helix-loop-helix (bHLH) domain, and extended MYC Box II (eMBII) come together to form a bivalent, high-affinity small-molecule MYC inhibitor (MYCi) binding site. CRISPR-tiling mutagenesis identified mutations in the vicinity of the eMBII and bHLH regions that together confer MYCi resistance. Importantly, acetylation of K148 in eMBII, which is essential for MYC oncogenicity in vivo, enhanced MYCi binding affinity and is predicted to increase the structural order of this region. Furthermore, MYCi selectively modulated the expression of the same genes regulated by lysine-acetylated MYC in cancer cells. These studies provide a rationale for selective targeting of acetylated, oncogenic MYC with small molecules.

---

Intrinsically disordered proteins (IDPs) and protein regions (IDRs) lack stable tertiary structures yet play critical roles in cellular signaling and disease (*1, 2*). Their conformational plasticity makes them challenging therapeutic targets, particularly in cancer, where IDPs like MYC are frequently dysregulated but remain undruggable due to the lack of structured binding pockets. The MYC oncoprotein is a paradigmatic IDP and a master regulator of cancer hallmarks, driving proliferation, metabolism, and immune evasion, making it a highly attractive therapeutic target (*3–5*). Unlike other oncoproteins such as RAS, MYC is generally not mutated in human cancer, making it difficult to selectively target its oncogenic form (*6*). However, post-translational modifications in cancer cells may influence MYC oncogenic activity. For example, acetylation at lysines 148 (149 in mouse) and 157 (158 in mouse) are required for MYC oncogenicity in vivo (*7*). This suggests that specific targeting of acetylated MYC in cancers could increase the therapeutic window for MYC inhibitors. Structurally, MYC consists of a largely disordered N-terminal region containing conserved transcriptional regulatory elements (MYC boxes I–IV; MBI–IV) followed by a conserved basic-helix-loop-helix leucine zipper (bHLH-LZ) domain that mediates dimerization with MAX and DNA binding. The absence of discernible “druggable” pockets in MYC notwithstanding, several studies have empirically identified small molecules that interact with MYC and block its interaction with MAX and/or DNA (*8, 9*). One such small molecule MYC inhibitor, MYCi975, selectively binds to MYC and has demonstrated preclinical efficacy and remarkable tolerability (*10–12*). Yet, the structural basis for this and other MYC inhibitors remains unclear. Attempts at answering this question have focused on studying small molecule binding to the DNA/MAX-interacting interface of MYC and have found that a majority interact with a short stretch of amino acids in the bHLH-LZ domain called the “MycHot” region (*8, 13*). Recently, the N-terminal transactivation domain (TAD) termed coreMYC has been shown to undergo a conformational switch upon low-affinity, non-specific binding of small molecules like epigallocatechin gallate (EGCG) (*14*). However, the binding affinities of molecules to this region are relatively modest (in the micromolar range), and a clear structural mechanism for how these molecules specifically inhibit oncogenic MYC activity is lacking.

While investigating how MYCi975 binds to MYC, we made the serendipitous finding that the bHLH-LZ domain cooperates with a previously unrecognized region overlapping the MBII domain (extended MBII or eMBII). The fact that acetylation of lysines K148/149 and K157/158 (human/mouse) found within the eMBII region is required for in vivo MYC oncogenicity further motivated us to investigate how these post-translational modifications affect small molecule binding.

## RESULTS

### Two separate regions in MYC form a bivalent site for small molecule binding

Using a competitive fluorescence polarization assay, we previously estimated the MYC inhibitor MYCi975 binds to the MYC bHLH-LZ domain with a K_d_ ∼ 2.5 µM (*10*). To obtain a direct K_d_ measurement with unmodified ligand and protein, we established a biolayer interferometry (BLI) assay with full-length recombinant MYC protein. We used a MYCi975 regioisomer, compound 733, which lacks MYC inhibitory activity, as a negative control. Surprisingly, MYCi975 bound full-length MYC with a K_d_ = 262 ± 3.8 nM, about an order of magnitude tighter than the estimated K_d_ for the bHLH-LZ domain only (Fig. 1A, S1A). These results are consistent with the possibility of an additional region of MYC contributing to MYCi975 binding. To test this hypothesis, we used a series of HA-tagged MYC mutant proteins with sequential tiled deletions of 63 amino acids spanning the entire sequence (Fig. 1B). We expressed these mutant proteins in HEK293 cells and used biotinylated-MYCi975 (*10*) to pull down bound proteins, followed by western blot analysis. Deletion of the bHLH-LZ MYCi-binding region (ΔG mutant, amino acids 379-439) only reduced binding by ∼50 percent, consistent with the presence of another MYCi975 binding site within MYC. Indeed, deletion of amino acids 127-189 (ΔC mutant) also reduced binding by about ∼50% (Fig. 1B). This region, which encompasses the conserved MBII domain of MYC (amino acids 128-143), has recently been reported to contain a “conformational switch” that cycles between a closed, inactive, and an open, active conformation, modulating interaction with MBII-binding proteins (*14*). We hypothesized that this part of MYC, spanning amino acids 127-189 (herein referred to as extended MBII or eMBII region) contributes to MYCi975 binding. To investigate this directly, we generated a series of recombinant mutant MYC proteins and tested them for binding to MYCi975 in the BLI assay. The binding affinity of MYCi975 for the bHLH-LZ domain was 2230 ± 3.4 nM, similar to the ∼2500 nM affinity estimated by the competitive fluorescence polarization method reported by Han et al (*10*). Deletion of the eMBII region from full-length MYC also drastically reduced MYCi975 affinity (K_d_ = 2100 ± 2.2 nM), a value that is similar to MYCi975 affinity to the isolated bHLH-LZ domain. A MYC mutant in which the bHLH-LZ region was deleted showed binding to MYCi975 but had rapid dissociation kinetics, precluding K_d_ determination (Fig. 1C).

**Figure 1.**
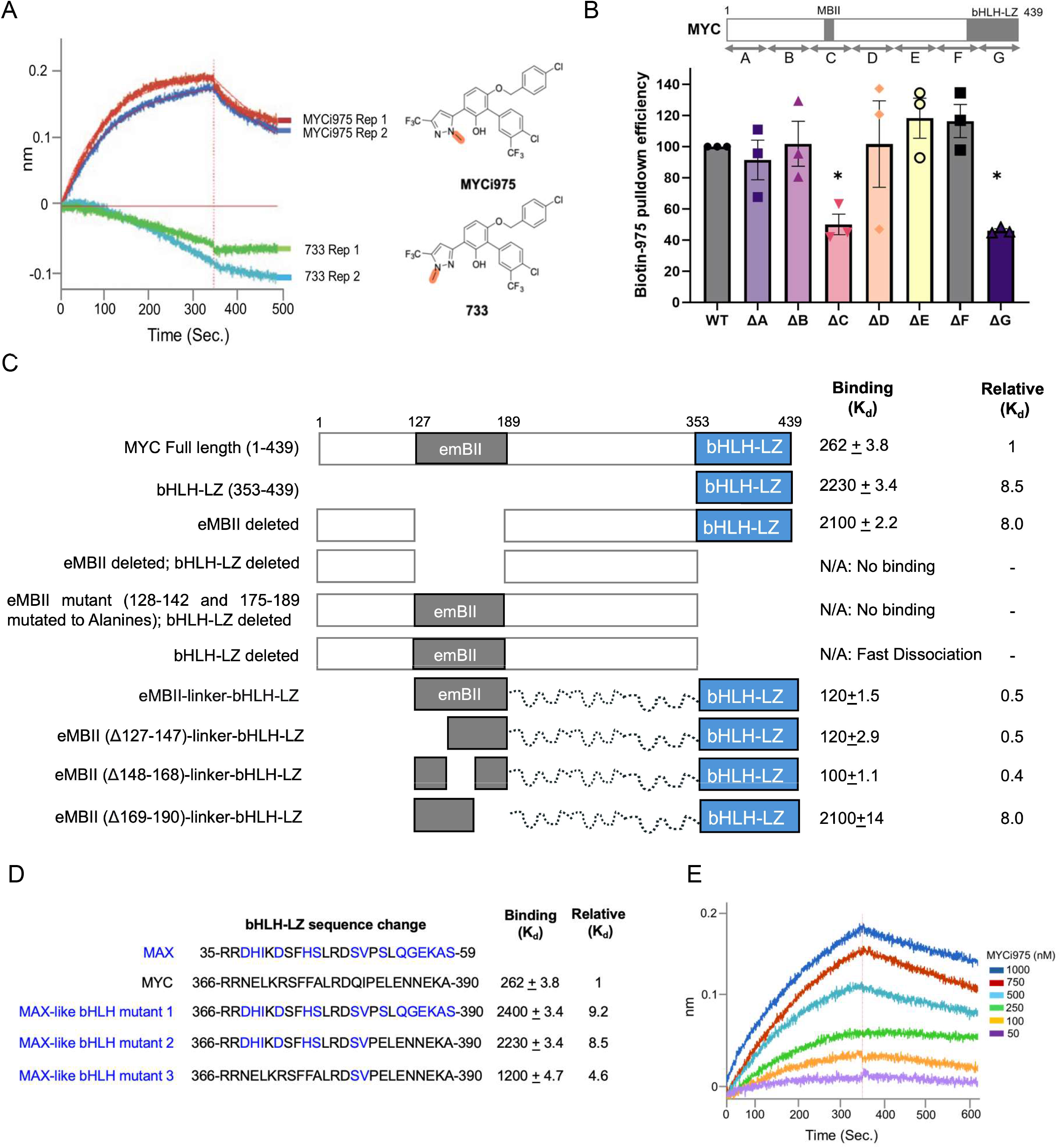
MYC contains a bivalent small molecule binding site. **(A)** Biolayer interferometry (BLI) sensorgrams showing association of MYCi975 and the negative control compound 733 with recombinant WT MYC. **(B)** Pulldown of HA-tagged WT MYC and MYC deletion mutants (ΔA to ΔG) using biotinylated MYCi975 (Biotin-975), followed by western blot. **(C)** Binding affinities (BLI K_d_) of MYCi975 for recombinant WT MYC and MYC mutants. **(D)** bHLH-LZ regions of MYC and MAX and the mutations generated. **(E)** BLI K_d_ of recombinant MYC eMBII linked to bHLH with MYCi975.

We have shown previously that MYCi975 does not bind to MAX (*10*) despite the overall similarity of the bHLH-LZ domains of the two proteins. We therefore generated three mutant MYC proteins in which the bHLH-LZ region was altered to mimic the MAX sequence (MAX-like bHLH mutants 1-3; Fig. 1D); we then used BLI to assess the relative importance of this region to MYCi975 binding in the context of the full-length MYC protein. In all the MYC mutants, the binding affinity of MYCi975 was reduced by 4.6- to 9-fold relative to wild-type MYC protein (K_d_ = 2400 ± 3.4 nM for mutant 1, 2200 ± 3.4 nM for mutant 2 and 1200 ± 4.7 nM for mutant 3) (Fig. 1D). Furthermore, deletion of the bHLH-LZ region concurrent with either deletion or alanine mutagenesis of the eMBII region completely abrogated MYCi975 binding (Fig. 1C). Notably, we generated a mutant in which the bHLH-LZ region was linked to the eMBII region with a flexible 40-amino acid glycine-rich linker. This mutant binds efficiently to MYCi975 (K_d_ = 120 ± 1.5 nM), approximately two-fold better than the wild-type MYC protein (Fig. 1C, 1E). We generated three additional bHLH-LZ-eMBII constructs containing progressive deletions within the eMBII region (residues 127–190). The first two deletion mutants retained MYCi975 binding affinity within approximately one order of magnitude of full-length MYC. However, the third mutant, carrying a deletion spanning residues 169-190, exhibited a drastically reduced MYCi975 binding affinity (K_d_ = 2100 + 14 nM), comparable to the affinity of MYCi975 for the isolated bHLH-LZ domain alone, suggesting that this sub-region of eMBII makes a critical contribution to MYCi975 binding (Fig. 1C). These results suggest that eMBII and bHLH-LZ probably create a cooperative MYCi975 binding pocket with bHLH-LZ as the primary driver domain for binding. Disorder prediction analysis (PONDR-VSL2) indicates that MYC contains two relatively ordered regions, the bHLH-LZ domain and eMBII, flanked by a highly disordered N-terminal transactivation domain (Fig. S1B). Together, these results support a bivalent MYCi binding model to MYC.

### Nuclear magnetic resonance spectroscopy and molecular docking models of bHLH-LZ, eMBII and MYCi975

To further characterize the molecular interactions driving small-molecule engagement, we collected 2D heteronuclear single quantum coherence (HSQC) NMR spectra using uniformly ^15^N-labeled bHLH-LZ. Addition of MYCi975 to bHLH-LZ induces chemical shift perturbations (CSPs) and modest intensity changes in a small number of resonances, relative to the apo bHLH-LZ baseline, indicative of a specific binding site. These perturbations reflect fast-intermediate exchange on the NMR timescale, which is consistent with the low micromolar binding affinity measured for the isolated domain (Fig. 1C, 2A). In contrast, incubation of bHLH-LZ with the eMBII fragment (Fig. 2B) produced widespread CSPs accompanied by noticeable resonance line-narrowing and peak sharpening across a distinct subset of backbone amides indicating complexation of these two regions. Crucially, when MYCi975 is added to the preformed bHLH-LZ: eMBII complex (Fig. 2C), the resulting spectrum contains minimal additional perturbations. This spectral similarity suggests that the inter-domain association between eMBII and bHLH-LZ remains the predominant conformational state in the presence of MYCi975. Together, these solution NMR results provide support for a model in which eMBII and bHLH-LZ cooperatively participate in small-molecule recognition within full-length MYC.

**Figure 2.**
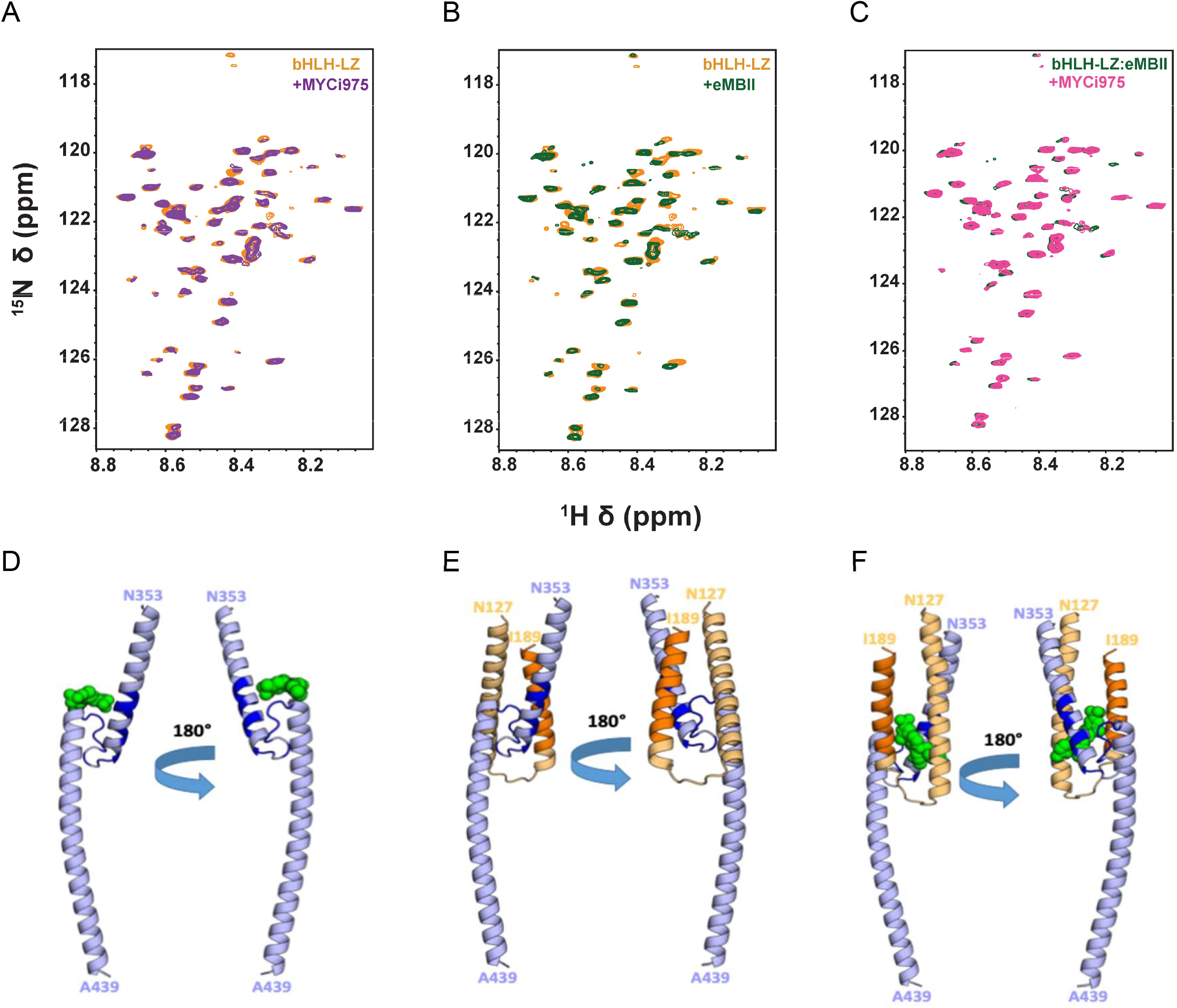
^1^H-^15^N HSQC spectra and molecular docking models of MYC bHLH-LZ, eMBII and eMBII/MYCi975 ternary complexes. **(A)** Overlay of the 2D ^1^H-^15^N HSQC spectrum of free bHLH (orange) with that of the bHLH: MYCi975 complex (violet). **(B)** Overlay of the 2D ^1^H-^15^N HSQC spectrum of free bHLH (orange) with that of the bHLH: eMBII complex (dark green). **(C)** Overlay of the 2D ^1^H-^15^N HSQC spectrum of the bHLH: eMBII complex (dark green) with that of the bHLH: eMBII: MYCi975 complex (pink). **(D)** Top scoring, unrestrained, HADDOCK docking models of bHLH-LZ: MYCi975 binary complex. **(E)** The docking model of bHLH-LZ: eMBII binary complex. **(F)** Critical interaction residues, previously identified via mutagenesis and biolayer interferometry (BLI), are highlighted in dark blue on the bHLH-LZ domain (light blue) and dark orange on the eMBII domain (light orange), respectively. MYCi975 is represented by a green spherical model.

To develop a molecular model for how the bHLH-LZ: eMBII domains complex with MYCi975, unrestrained molecular docking calculations were performed using HADDOCK (*15*). The best scoring poses (Fig. 2D, E, F) show that the predicted bHLH-LZ: eMBII binary interface (Fig. 2E) coincides with the key interaction residues identified by BLI (Fig. 1), while the experimentally docked MYCi975 inhibitor (Fig. 2D) is positioned at the same natural protein-protein interface. The highest scoring ternary model (Fig. 2F) shows MYCi975 situated directly between the bHLH-LZ and eMBII domains with simultaneous binding to the key binding surfaces of both proteins.

### *MYC* CRISPR-tiling mutagenesis screen supports a bivalent MYCi binding model and identifies resistance alleles

In parallel studies, we had undertaken an unbiased CRISPR-tiling mutagenesis screen (*16*) with the goal of identifying potential *MYC* alleles that lead to resistance to MYCi975. This is the gold standard approach for demonstrating on-target activity for a small molecule. Since *MYC* is essential for the viability of cancer cells, this screen faces the added challenge that any induced mutations must retain sufficient MYC function for cell viability to score positively in this assay. We generated a single guide (sg) RNA tiling library consisting of 886 guides covering the entire *c-MYC* gene and 114 control guides. The sgRNA library was introduced into Cas9-expressing PC3M cells (contain 5-6 copies of the *MYC* gene; Fig. S2A) and the cells treated with vehicle or 6µM MYCi975 for up to 35 days (Fig. 3A). We sequenced the cells at baseline and after 21 and 35 days of treatment and assessed guide enrichment/depletion. The entire screen was performed independently by two different investigators. Notably, guides enriched in MYCi975-treated cells did not cluster in any single region of MYC, consistent with our observation that more than one part of the MYC protein contributes to overall MYCi975 binding (Fig. 1C). Three guides were enriched in both screens, overlapping the eMBII region (A169) or the bHLH-LZ domain (A361 and A423) (Fig. 3B). To validate these observations, we introduced pairs of guides from these two regions (i.e. A169+A361 or A169+A423) into PC3M-Cas9 cells followed by MYCi975 treatment; strikingly, cells transfected with the combination of guides were partially resistant to MYCi975 (Fig. S2B). By contrast, cells co-transfected with a guide that was not reproducibly enriched in the screens (A11), failed to survive MYCi975 treatment (Fig. S2C). To define the genetic basis of resistance to MYCi975, we performed deep sequencing of the MYC locus in resistant cell populations, identifying mutations that non-randomly clustered in the vicinity of the eMBII and bHLH-LZ regions (Fig. 3B, S2D, E), consistent with the dynamic interaction model of small molecules with IDPs (*17–19*). It is notable that no alterations were observed in any of the conserved residues critical for DNA binding or leucine-zipper mediated dimerization with MAX in the bHLH-LZ domain.

**Figure 3.**
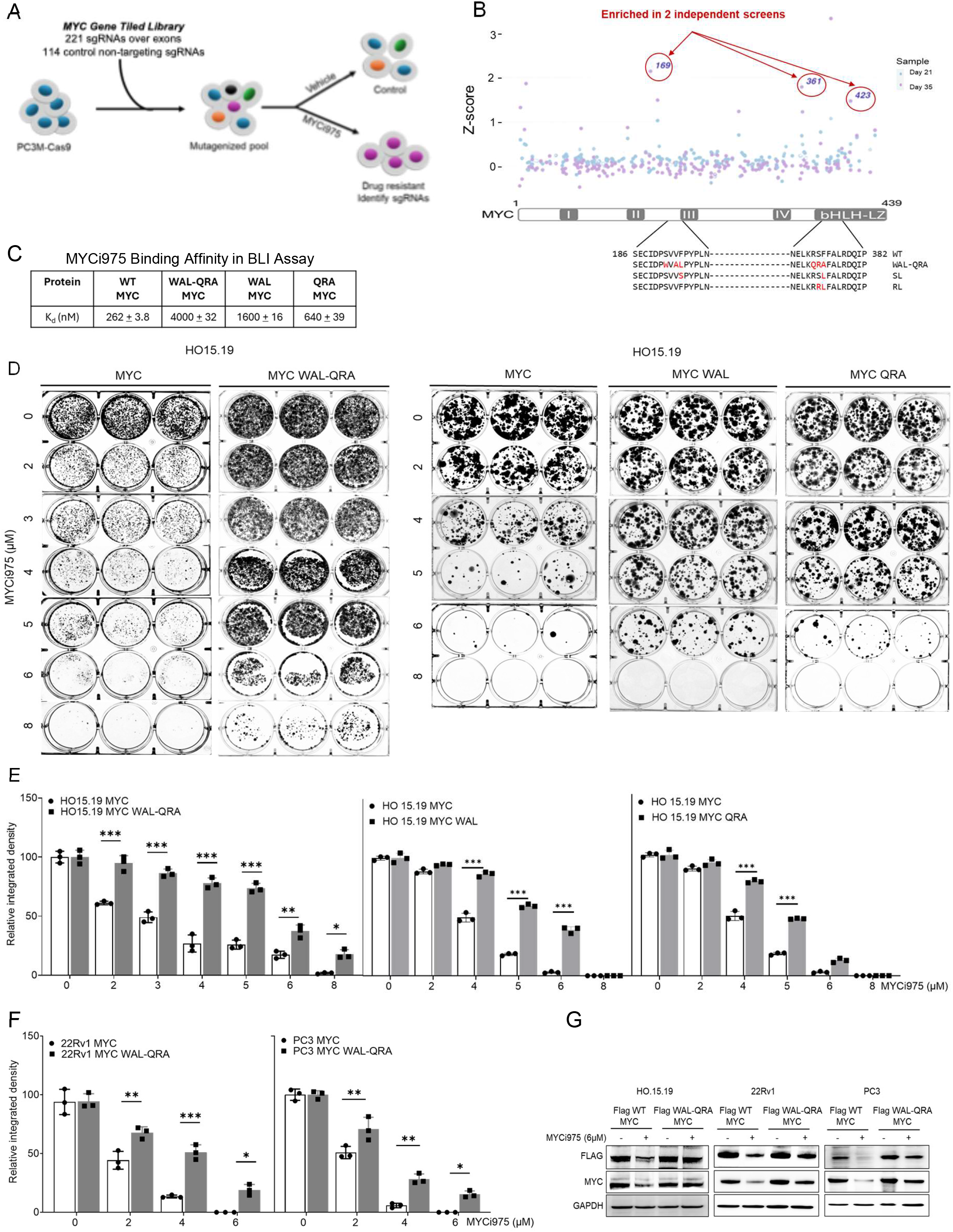
CRISPR tiling screen identifies functional MYC regions critical for MYCi975 sensitivity. **(A)** Schematic of the CRISPR Cas9 suppressor screening strategy. A Cas9-expressing PC3M cell pool was transduced with a pooled sgRNA library targeting the MYC locus. Following enrichment in 6 µM MYCi975, resistant clones were isolated and sequenced to identify mutations conferring MYCi975 resistance. **(B)** Volcano plot summarizing the enrichment of sgRNAs guides across two independent screens. sgRNA guides targeting residues A169, A361, and A423 were significantly enriched. Deep sequencing of resistant clones identified mutation hotspots clustering close to the eMBII region (amino acid residues 192–195) and the bHLH (amino acid residues 372-374) domains of MYC. Representative sequences are shown. **(C)** BLI K_d_ of MYCi975 for recombinant WT MYC, WAL-QRA MYC mutants, and individual domain MYC mutants (WAL and QRA). **(D)** Colony formation assay of HO15.19 fibroblast cells stably expressing Flag-tagged WT MYC, WAL-QRA, WAL and QRA MYC after treatment with DMSO and varying concentrations of MYCi975 for 14 days. Representative wells of colony formation assay are shown. **(E)** Quantification of colony formation assays in HO15.19 fibroblast cells expressing Flag-tagged WT MYC, WAL-QRA, WAL and QRA MYC mutants after MYCi975 treatment for 14 days. Data was normalized to DMSO control and represented the mean ± S.D. from biological replicates. *Denotes statistical significance (*p<0.05; **p<0.01; ***p<0.001). **(F)** Quantification of colony formation assays in 22Rv1 and PC3 cells expressing Flag-tagged WT MYC and WAL-QRA MYC mutants after MYCi975 treatment for 14 days. Data was normalized to DMSO control and represented the mean ± S.D. from biological replicates. *Denotes statistical significance (*p<0.05; **p<0.01; ***p<0.001). **(G)** Immunoblot analyses of FLAG-tagged MYC protein stability in HO15.19, 22Rv1 and PC3 cells expressing Flag-MYC WT or Flag-WAL QRA-MYC mutant, treated with 6µM MYCi975 for 24 hours. GAPDH serves as a loading control.

To directly test the sufficiency of the identified mutations in promoting resistance to MYCi975 treatment, we generated a plasmid encoding one of the mutant *MYC* alleles identified: three alterations near the eMBII region (S192W, V193V, V194A, F195L, abbreviated and referred to as WAL) and three in the bHLH-LZ domain (R372Q, S373R, F374A, abbreviated and referred to as QRA). First, we determined the impact of mutations on MYCi binding. We expressed recombinant MYC containing the WAL-QRA, WAL or QRA alone mutations and assessed MYCi975 binding affinity in the BLI assay. MYCi975 bound poorly to the WAL-QRA MYC mutant, with K_d_ of 4µM (Fig. 3C, S2F), ∼15-fold weaker than recombinant wild-type MYC. Consistent with our bivalent binding model, the WAL and QRA MYC mutants showed intermediate binding affinities of 1600 nM and 640 nM respectively. We next generated stable cell lines expressing Flag-tagged WT-MYC, WAL-QRA, WAL, and QRA MYC proteins. Stable expression of the WAL-QRA MYC mutant in HO15.19 *MYC*-null fibroblasts, as well as in 22Rv1 and PC3 prostate cancer cells, conferred increased resistance to MYCi975 treatment compared to cells expressing WT-MYC, WAL or QRA-MYC alone (Fig. 3D, E, F, S2G, S2H). Furthermore, MYCi975-induced degradation of WAL-QRA MYC mutant was reduced compared to wild-type MYC and WAL or QRA-MYC alone (Fig. 3G, S2I, S2J) consistent with MYCi binding-dependent degradation of MYC (*10*). The results indicate that the WAL-QRA mutations promote resistance by reducing MYCi binding to MYC.

### Lysine acetylation in the eMBII region modulates MYCi binding affinity

We noted the presence of lysine residues K148/149 and K157/158 (in human/mouse respectively) in the eMBII region. Acetylation of these lysine residues (e.g. by p300/CBP) has been observed in cancer and promotes MYC oncogenic activity by regulating specific MYC target gene programs (*7, 20–22*). To assess whether and how K148/K157 acetylation may affect MYCi binding, we first evaluated the impact of acetylation on the predicted structure of this region in AlphaFold 3 (*23*). The models indicate that K148/K157 acetylation increased the stability of the helix containing these lysine residues (Fig. 4A, B, S3A). To investigate whether this change may affect MYCi-binding, we generated MYC mutants in which K148/K157 were mutated to glutamines (Q) as an acetylation mimic or unacetylatable arginine (R) residues. We generated recombinant K148Q/K157Q mutant MYC to test MYCi975 binding in the BLI assay (Fig. S3B). The MYC K148R/K157R mutant was used as a control. MYCi975 bound to the K148R/K157R mutant with a similar affinity as wild-type MYC (K_d_ = 230 ± 6.1 nM vs 262 ± 3.8 nM). By contrast, the K148Q/K157Q mutant showed a higher-affinity binding (K_d_ = 130 ± 3.8 nM). These results suggest that acetylated, oncogenic MYC may bind better to MYCi975. Although the K-to-Q mutation mimics the loss of positive charge of the acetyllysine, it does not completely recapitulate the changes induced by lysine acetylation. We therefore generated recombinant MYC with specifically acetylated lysine residues at positions 148 and 157 by genetically encoding acetyllysine (AcK) in Escherichia coli (E. coli) (*24, 25*). The purified protein was characterized by mass spectrometry to confirm acetylation of K148 and K157 and by probing with AcK148-MYC specific antibodies (*7*) (Fig. 4C, S3C). Using the BLI assay, we found that the binding affinity of MYCi975 to the AcK148/AcK157 MYC mutant was increased relative to the wild-type MYC protein (K_d_ = 90 ± 1.1 nM vs 262 ± 3.8 nM) (Fig. 4D, S3D). Next, we examined whether MYCi975 preferentially binds to acetylated MYC in 22Rv1 prostate cancer cells. To this end, we used biotinylated-MYCi975 (*10*) to pull down MYC from 22Rv1 and examined the AcK148-MYC acetylation status of the recovered protein by probing with AcK148-MYC specific antibodies (*7*). Biotin-MYCi975 recovered higher levels of AcK148-MYC relative to total MYC, suggesting that in cells, MYCi975 has a higher affinity for more oncogenic AcK148-MYC compared to non-acetylated MYC (Fig. 4E, F).

**Figure 4.**
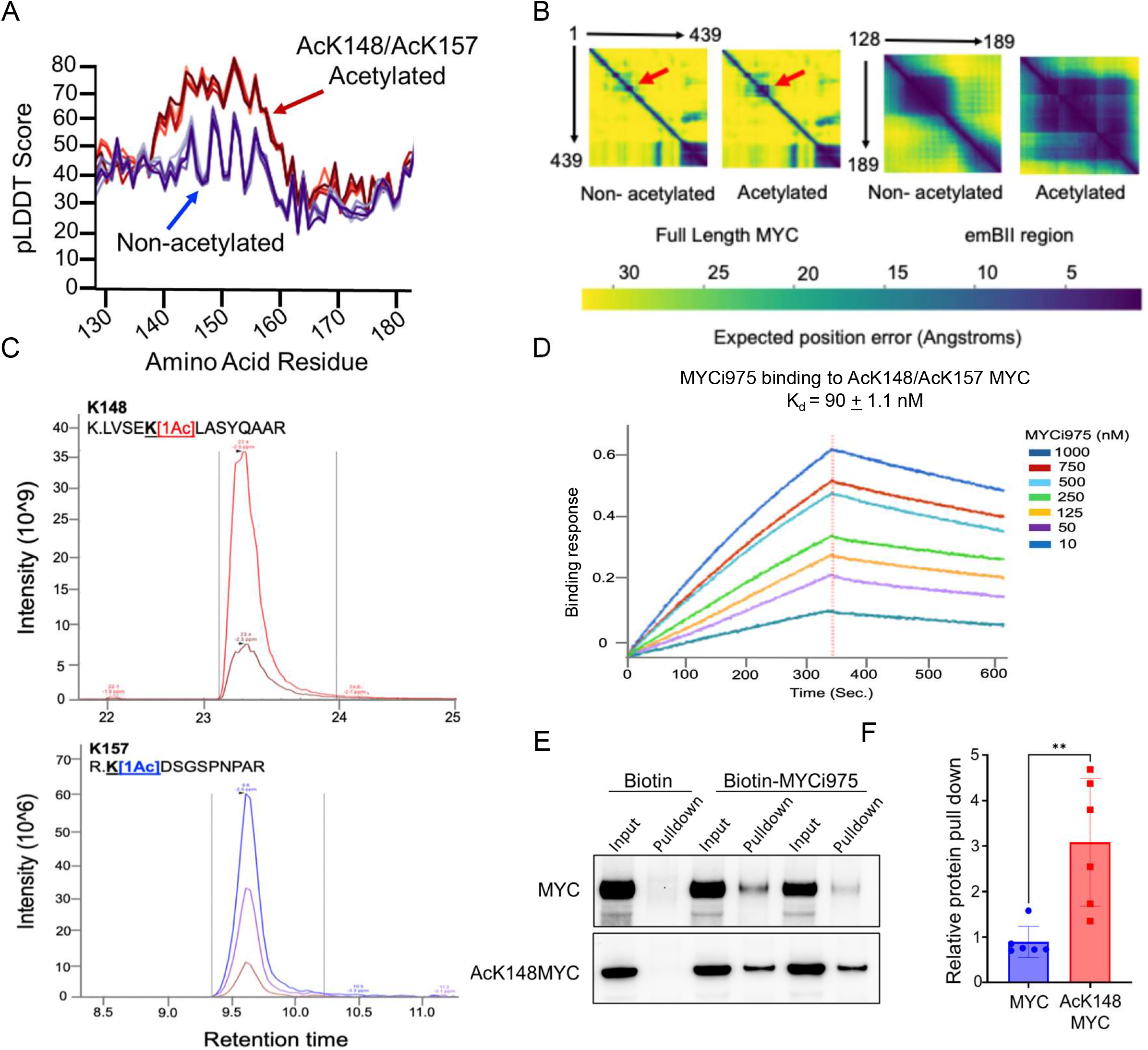
Acetylation of MYC lysine-148 and lysine-157 stabilizes the eMBII region and enhances MYCi975 binding. **(A)** AlphaFold 3 predicted local distance difference test (pLDDT) scores for unacetylated (blue) and acetylated (red) MYC models, highlighting increased structural stability upon K148/K157 acetylation. **(B)** Predicted aligned error (PAE) maps for full-length MYC (residues 1–439) and for the extended eMBII region (residues 128–189) in unacetylated and acetylated states. **(C)** Mass spectrometric confirmation of site-specific acetylation at K148 and K157 in purified recombinant MYC expressed by genetically encoding acetyllysine (AcK) in Escherichia coli (E. coli). **(D)** BLI K_d_ of MYCi975 to acetylated MYC (AcK148/AcK157-MYC). **(E)** Pulldown of endogenous MYC and AcK148-MYC using biotinylated MYCi975 (Biotin-975), followed by immunoblotting with total MYC (Y69) and AcK148-MYC specific antibody. **(F)** Quantification of pull-down efficiency for MYC and AcK148-MYC from n=6 independent experiments (p<0.01). *Denotes statistical significance (*p<0.05; **p<0.01; ***p<0.001).

### Pan-cancer acetylome analysis reveals tumor-enriched acetylation of MYC at K148 across human cancers and is preferentially targeted by MYCi975

We analyzed high-resolution acetylome data from the Clinical Proteomic Tumor Analysis Consortium (CPTAC) to assess the abundance of MYC acetylation at residue K148. We analyzed 8 cohorts, totaling 928 tumors and 389 normal tissues. MYC acetylation was significantly elevated in tumors compared to matched solid normal tissues in the majority of cohorts examined (5 of 8), including breast cancer (PDC000239; p<0.0001), lung squamous cell carcinoma (PDC000233; p<0.0001), lung adenocarcinoma (PDC000491; p=0.0184), glioblastoma (PDC000245; p=0.028) and multiple tumors (PDC000450; p=0.009) (Fig. 5A). One lung cancer cohort (PDC000224) and the two uterine corpus endometrial carcinoma cohorts (PDC000226, PDC000443) did not follow this pattern.

**Figure 5.**
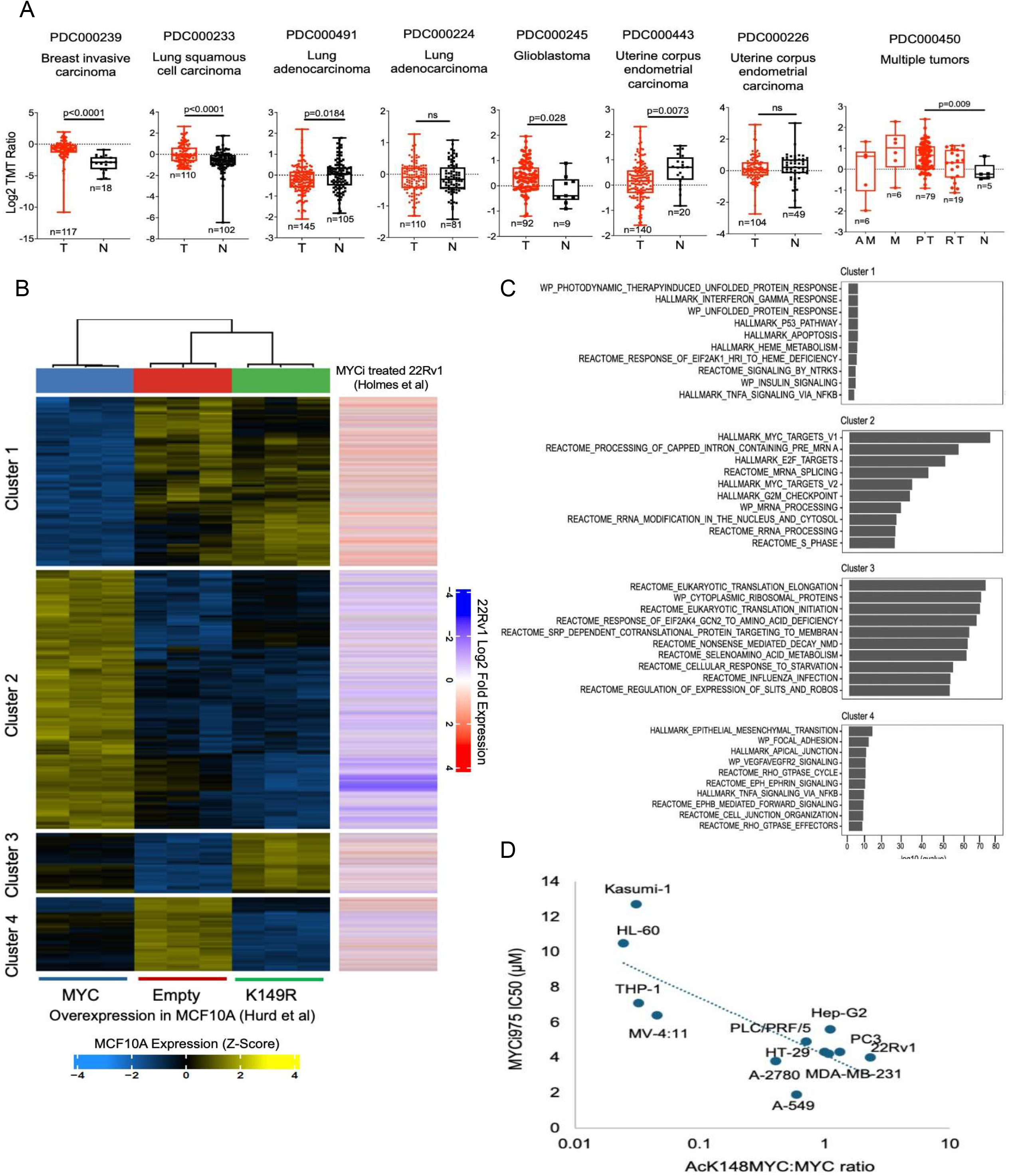
Pan-cancer acetylome analysis reveals tumor-enriched AcK148-MYC and MYCi975 selectively inhibits AcK148-MYC activity. **(A)** We analyzed high-resolution acetylome data from the Clinical Proteomic Tumor Analysis Consortium (CPTAC), including 8 cohorts totaling 928 tumors and 389 normal tissues. K148 MYC modification was significantly elevated in tumors compared to matched solid normal tissues in 5 of 8 cohorts examined: breast cancer (PDC000239; p<0.0001), lung squamous cell carcinoma (PDC000233; p<0.0001), lung adenocarcinoma (PDC000491; p=0.0184), glioblastoma (PDC000245; p=0.028), and a multi-cancer cohort (PDC000450; p=0.009) comprising breast cancer, epithelial and complex epithelial neoplasms, glioblastoma, lung cancer, meningiomas, skin cancer and uterine adenocarcinoma. One lung cancer cohort (PDC000224) and two uterine corpus endometrial carcinoma cohorts (PDC000226, PDC000443) did not follow this pattern. Each data point represents an individual patient’s sample. Statistical analysis was performed using a two-tailed Mann-Whitney U test. p-values: ****p<0.0001, *p<0.05. (Abbreviations: AM, Additional Metastatic; M, Metastatic; PT, Primary Tumor; RT, Recurrent Tumor; N, Normal). **(B)** Unsupervised clustering of MYC target genes based on responses to wild-type MYC versus acetylation-deficient MYC (K149R) identified four clusters comprising MYC-repressed (clusters 1, 4) and MYC-induced (clusters 2, 3) programs. MYCi975 selectively induced acetylation-dependent stress-response genes (cluster 1) and repressed acetylation-dependent canonical MYC targets (cluster 2), while minimally affecting acetylation-independent programs (clusters 3, 4), indicating preferential disruption of acetylated MYC dependent transcription. **(C)** Functional enrichment analysis of each gene cluster highlighting the top 10 significantly enriched pathways. **(D)** Inverse correlation between AcK148-MYC abundance and sensitivity to MYCi975. Scatter plot showing the inverse relationship between the relative abundance of the AcK148-MYC modification (AcK148-MYC:MYC ratio) and cellular sensitivity to the MYC inhibitor MYCi975 (IC50) across a panel of 12 cancer cell lines including leukemia (Kasumi-1, HL-60, THP-1, MV-4:11), Prostate (PC3 and 22Rv1), Liver (PLC/PRF/5, HepG2), lung (A549), breast (MDA-MB-231), ovarian (A2780) and colorectal (HT-29) cell lines. A higher ratio of AcK148-MYC is associated with a lower IC50, indicating increased sensitivity to MYCi975.

A previous study identified a subset of MYC target genes that are selectively regulated by acetylated MYC in a model of MYC-transformed mammary epithelial MCF10A cells (*7*) by overexpressing mouse MYC or K149R (K148 in human) mutant. To determine if these “acetylated-MYC target transcriptional genes” are selectively targeted by MYCi975 treatment, we reanalyzed the published data sets from Hurd et al (*7*) and our own data published in Holmes et al (*11*). Unsupervised clustering of MYC target genes based on their response to wild-type MYC versus the acetylation-deficient MYC K149R mutant identified four distinct gene clusters (Fig. 5B). Clusters 1 and 4 comprised MYC-repressed gene programs, whereas clusters 2 and 3 represented MYC-induced targets. Pathway enrichment analysis revealed that cluster 1 was enriched for unfolded protein response, interferon signaling, and stress-associated pathways, while cluster 4 was enriched for epithelial to mesenchymal transition (EMT), focal adhesion, and cell to cell junction pathways. In contrast, cluster 2 was enriched for RNA processing, MYC target genes, and cell cycle progression, while cluster 3 was enriched in pathways involved in eukaryotic translation and oxidative phosphorylation (Fig. 5C).

Strikingly, only clusters 1 and 2 exhibited a strict dependence on MYC acetylation, as expression of the K149R mutant abrogated MYC-mediated repression (cluster 1) or activation (cluster 2). In contrast, regulation of clusters 3 and 4 remained largely intact in the presence of K to R mutation, indicating that these transcriptional programs are acetylation independent. Notably, MYCi975 treatment resulted in robust induction of the acetylation-dependent cluster 1 genes and selective repression of cluster 2 genes, while exerting relatively modest effects on acetylation-independent clusters (Fig. 5B). Notably, cluster 2 genes include MYC-induced programs critical for cell cycle progression and metabolic activity, consistent with prior reports that these genes are enriched for low-affinity MYC binding sites and are particularly sensitive to perturbations in MYC transcriptional activity (*26*). Together, these data indicate that MYCi975 preferentially disrupts AcK148-MYC dependent transcriptional program, selectively suppressing MYC-driven cell cycle and biosynthetic programs while derepressing stress response pathways. Finally, we investigated the response to MYCi975 in a panel of 12 diverse cancer cell lines to assess a possible link between MYCi975 sensitivity and AcK148-MYC levels. We found an inverse correlation between the AcK148-MYC:MYC ratio and MYCi975 IC_50_ in these cells (Fig. 5D, S4A).

### MYCi648, a close analog of MYCi975, exhibits improved MYC binding and antitumor activity

To test the bivalent binding model beyond MYCi975, we tested additional MYC inhibitors that disrupt MYC/MAX interaction, including the previously described inhibitor MYCi361 (*10*). We also took advantage of compounds identified in a medicinal chemistry campaign we conducted to develop more potent analogs of MYCi975. Compounds were tested in a MYC-dependent E-box luciferase reporter assay, a control CMV-luciferase reporter assay, as well as cytotoxicity to MYC-dependent PC3 cells and control MYC-independent PC12 cells. Of 105 MYCi analogs synthesized and tested, MYCi648 (NUCC-0202648) (Fig. 6A) was the most potent and selective in the reporter and cytotoxicity assays (Fig. 6B). Both MYCi361 and MYCi648 showed preferential higher binding to acetylated MYC relative to non-acetylated MYC protein (Fig. 6B).

**Figure 6.**
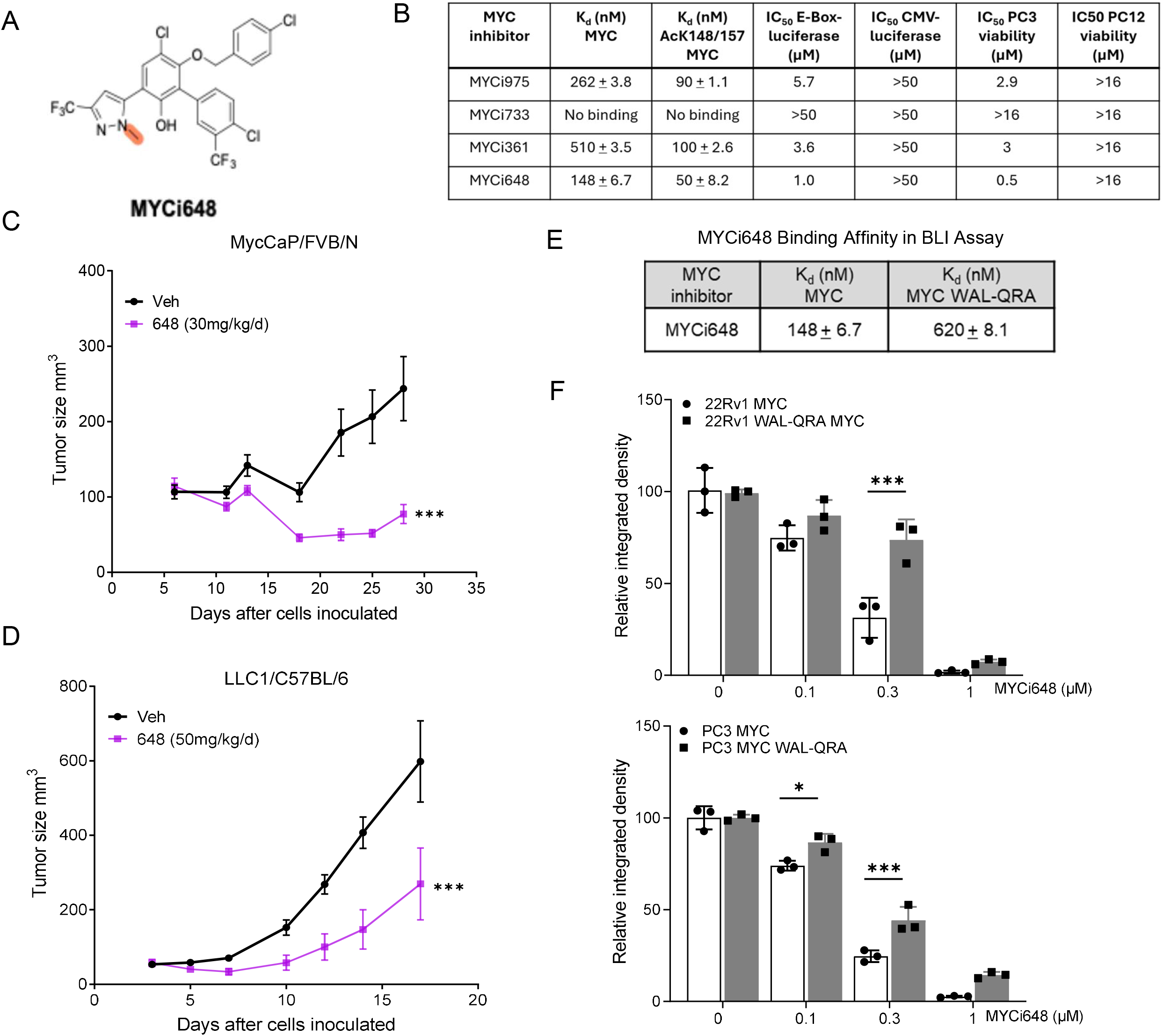
MYCi648 is a potent MYC inhibitor with improved antitumor efficacy. **(A)** Chemical structure of the new MYCi analog MYCi648. **(B)** Binding affinities K_d_ of MYC inhibitors to WT MYC and acetylated MYC (AcK148/157-MYC), along with cellular activity measured by E-box luciferase inhibition, CMV luciferase control assays, and viability assays in PC3 and control PC12 cells. **(C)** Tumor growth inhibition (TGI) in the MycCaP/FVB/N prostate cancer model treated with vehicle (n=8) or MYCi648 (30 mg/kg/d; n=5) i.p. (***p<0.001). **(D)** Tumor growth inhibition in the LLC1 syngeneic model (C57BL/6 mice) treated with vehicle (n=6) and MYCi648 (50 mg/kg/d; n=6) i.p. (***p<0.001). **(E)** BLI K_d_ of MYCi648 to WT MYC and the WAL-QRA MYC mutant, demonstrating reduced binding to the WAL-QRA MYC mutant protein. **(F)** Quantification of colony formation assays in 22Rv1 and PC3 cells expressing WT MYC and WAL-QRA MYC mutants after MYCi648 treatment for 14 days. Data was normalized to DMSO control and represented the mean ± S.D. from biological replicates. *Denotes statistical significance (*p<0.05; **p<0.01; ***p<0.001).

Pharmacokinetic analysis at an oral dose of 1 mg/kg revealed that MYCi648 exhibited intrinsically lower systemic exposure than MYCi975. Dose-normalized AUC values were 1067 and 2556 ng-h/mL per mg/kg for MYCi648 and MYCi975, respectively, corresponding to a 2.4-fold difference. To compare absolute systemic exposure at the respective efficacy doses, AUC was extrapolated based on dose proportionality. At 50 mg/kg, the estimated absolute AUC for MYCi648 was 53,350 ng·h/mL. At 100 mg/kg, the estimated absolute AUC for MYCi975 was 255,600 ng·h/mL. Thus, the absolute systemic exposure to MYCi975 was 4.8-fold higher than that to MYCi648 at the doses used in the efficacy studies (Fig. S5A, B). Nevertheless, despite this, MYCi648 showed greater efficacy in vivo than MYCi975 in two immunocompetent syngeneic tumor models: MycCaP prostate tumors in FVB mice and LLC1 lung tumors in C57BL/6 mice. MYCi648 dosed at 30 mg/kg/d induced tumor regression (TGI >100%), compared to 52% TGI achieved by 100 mg/kg/d of MYCi975 (p<0.001) reported in (*10*) in the MycCaP model (Fig. 6C). In the LLC1 model (C57BL/6 mice), MYCi648 (50 mg/kg/d) achieved a TGI of 61.1%, nearly double the 32.5% TGI seen with the higher 100 mg/kg/d dose of MYCi975 (p<0.01) previously reported in (*10*) (Fig. 6D).

To examine whether the WAL-QRA mutation that impairs MYCi975 binding also affects the binding of MYCi648 to MYC, we performed the BLI binding assays. Binding of MYCi648 to the WAL-QRA MYC mutant was reduced 4-fold relative to wild-type MYC (K_d_ = 620 ± 8.1 nM vs 148 ± 6.7 nM) (Fig. 6E, S5C). Colony formation assays in 22Rv1 and PC3 cells showed partial resistance to MYCi648 in cells expressing the WAL-QRA MYC mutant (Fig. 6F). These studies indicate cross-resistance between MYCi648 and MYCi975 and support the bivalent binding model beyond MYCi975.

## Discussion

The results presented here address two major obstacles in targeting a paradigmatic IDP, the oncoprotein MYC: i) the lack of “druggable” pockets and ii) the absence of cancer-specific mutations that can be selectively targeted with small molecules. We show that two parts of MYC, the C-terminal bHLH-LZ domain and the eMBII region, together form a high-affinity binding small molecule binding site and NMR spectroscopy validated this model. MYCi975 induced localized chemical shift perturbations (CSPs) defining a discrete bHLH-LZ epitope, excluding nonspecific interactions. eMBII alone produced widespread CSPs with line-narrowing, indicating restricted bHLH-LZ dynamics. Critically, MYCi975 addition to the preformed bHLH-LZ:eMBII complex yielded minimal additional CSPs, with spectra superimposable on the complex alone. This establishes eMBII as the dominant conformational determinant and MYCi975 as engaging this composite surface. Collectively, these data confirm that eMBII and bHLH-LZ form a contiguous binding interface, coupling small-molecule recognition to a native intramolecular environment. Acetylation of lysines K148 and K157 in the eMBII region, essential for MYC oncogenic activity in vivo (*7*), increases the binding affinity of small molecule inhibitors to MYC. Notably, human tumors demonstrate significantly higher levels of acetylated MYC compared to normal tissues. Thus, preferential binding of small molecule inhibitors to “oncogenic” acetylated MYC provides a molecular basis for selective targeting of MYC in tumors. These new insights could inform future efforts for developing inhibitors that selectively bind to “oncogenic” acetylated MYC as supported by our findings with the better acetylated-MYC binder, MYCi648.

The bivalent binding model is strongly supported by orthogonal results from a CRISPR-tiling mutagenesis screen as resistance mutations were identified in both the eMBII and bHLH-LZ domains. While previous studies of MYC have identified small molecule interactions with the bHLH-LZ domain (*8*), the eMBII region has not been the focus of drug development efforts. The potential significance of the eMBII is suggested by a recent study that identified a conformational switch in this region that cycles between a closed, inactive, and an open, active conformational state and which can be modulated by the polyphenol epigallocatechin gallate (EGCG) (*14*). Based on the affinities of MYCi to the isolated bHLH-LZ and eMBII regions, we propose a binding model in which MYCi initially interacts with the bHLH-LZ region and the eMBII region is then recruited to stabilize this interaction. In this model, K148/K157 acetylation of the eMBII enhances this function.

The fact that K148/K157 acetylation enhances MYCi binding is fortuitous. A fundamental regulatory mechanism where lysine acetylation enhances MYC transcriptional activity and drives oncogenic programs (*7, 20, 22*) also simultaneously creates a therapeutic vulnerability by enhancing MYCi binding and resetting of the oncogenic MYC transcriptional program. As tumors generally express higher levels of AcK148-MYC, acetylation-dependent binding could possibly improve the therapeutic index of MYCi. Previous studies have shown that MYCi975 was well tolerated in mice (*10*). The preferential engagement of MYCi975 with acetylated-MYC may contribute significantly to this therapeutic window. Functionally, MYCi975 treatment selectively modulates acetyl-MYC dependent gene clusters, including cell cycle regulators, highlighting the functional relevance of K148 acetylation in sustaining the oncogenic MYC state and mediating therapeutic resistance. Supporting this, a recent study showed that in KRAS^G12D^ mutant pancreatic cancer, HDAC5 loss causes hyperacetylation of MYC at K148, which blocks NEDD4-mediated ubiquitination and degradation of MYC (*27*). Stabilized MYC protein maintains MAPK signaling and drives intrinsic resistance to KRAS inhibitors. Together, these findings suggest that tumors exploiting HDAC5 loss to resist KRAS-targeted therapies may, paradoxically, be sensitized to direct inhibition of acetylated MYC by MYCi.

Our findings align with emerging paradigms in protein acetylation and IDP regulation. Acetylation of IDRs has been shown to modulate liquid-liquid phase separation and membrane-less organelle formation, suggesting that MYC eMBII acetylation could similarly regulate its interactions and function (*28*). Moreover, K148 acetylation is intricately regulated by other post-translational modifications: TRAF6-mediated ubiquitination at K148 antagonizes acetylation and represses MYC’s oncogenic activity in myeloid malignancies, highlighting this residue as a critical regulatory node (*29*). Recent work also shows that acetate metabolism promotes K148 acetylation, stabilizing MYC and upregulating PD-L1, thereby linking MYC acetylation to tumor immune evasion (*30*). These observations suggest that tumors with dysregulated acetate metabolism may be particularly sensitive to MYCi therapies targeting acetylated MYC.

Our findings on MYC raise the possibility that other IDPs may be targeted through similar cooperative interactions between disordered domains, effectively “trapping” a specific conformation. Broadly, our results suggest the need to consider inter-molecular interactions and cancer-specific post-translational modifications when targeting oncogenic IDP proteins including MYC.

## Materials and Methods

### Mice

All animal experiments were approved by the Northwestern University Institutional Animal Care and Use Committee (IACUC) and conducted in accordance with ethical guidelines. Male FVB (for MycCaP model) and female C57BL/6 (for LLC1 model) mice (Jackson Laboratory) were housed in a specific pathogen-free facility and acclimated for at least 4 days prior to studies were initiated at 6–8 weeks of age. For the MycCaP allograft model, male FVB/N mice were injected subcutaneously with 1 × 10□MycCaP Ebox-Luc cells in 1:1 Matrigel/PBS. Tumor volume was calculated as (length × width²)/2. Upon reaching ∼100–120 mm³, mice were randomized to receive intraperitoneal MYCi648 (30 mg/kg/d) formulated in PBS with 10% DMSO, 20% Tween-80 or vehicle. For the LLC1 allograft model, female C57BL/6 mice were injected subcutaneously with 1 × 10□ LLC1 cells in 1:1 Matrigel/PBS. Three days post-implantation, mice were randomized to receive intraperitoneal MYCi648 (50 mg/kg/d) or vehicle.

### Cell culture

22Rv1, PC3, MycCaP, HEK293, MV-4:11, PC12, A549, LLC1 and HL-60 cell lines were purchased from the American Type Culture Collection (ATCC). TGR-1 and HO15.19 Rat-1 cells were from Professor John Sedivy (Brown University). THP-1 and Kasumi-1 lines were kindly provided by the laboratory of Dr. Yan Liu (Northwestern University). HT-29, HepG2, MDA-MB-231, A2780 and PLC/PRF/5 cell lines were kindly provided by the laboratory of Dr. Marcus Peter (Northwestern University) and PC3M by Dr. Matthew Clutter (Northwestern University). 22Rv1, PC3, PC3M, THP-1, HL-60, MDA-MB-231, and HT-29 cell lines were cultured in RPMI-1640 medium (Gibco, 11875-093). HEK293, LLC1, A2780, and MycCaP cells were cultured in DMEM (Gibco, 11965-118). HepG2 and PLC/PRF/5 cells were cultured in EMEM (ATCC, 30-2003), while MV-4:11 and Kasumi-1 cells were cultured in IMDM (ATCC, 30-2005). PC12 cells were grown in F-12K Medium (ATCC, 30-2004) with 2% fetal bovine serum (FBS; Gibco, A5256701) and 12.5% horse serum (Thermo Fisher Scientific, 16050122). All media were supplemented with 10% FBS and 1% penicillin-streptomycin (10,000 U/mL; Life Technologies, 15140-122). Cell culture was performed at 37°C and 5% CO_2_. Cell lines were tested for Mycoplasma using MycoAlert Mycoplasma Detection kit (Lonza, LT07-418).

### Biolayer Interferometry (BLI) assay

BLI binding experiments were performed using an Octet K2 system (ForteBio/Sartorius) in a solid black 96-well microplate at 37°C. Full-length MYC (UniProt P01106-1), MYC mutants and acetylated-MYC proteins were expressed as His-tagged proteins and purified. To determine the optimal ligand density for binding kinetic measurements, a loading optimization step was first performed by titrating the MYC proteins over the range of (1-25 µg/mL). Based on the optimization, a loading concentration of 20 µg/mL in binding buffer (50 mM HEPES (pH 7.4), 150 mM NaCl, and 10% DMSO; Table S1) was selected to achieve a consistent immobilization signal of 4–5 nM Ni-NTA biosensors (Sartorius). This loading level was chosen to provide a robust association signal for the highest analyte concentration while minimizing potential non-specific binding artifacts. The loading step was carried out for 3 minutes, followed by a 6-minute association phase and a 5-minute dissociation phase. MYC inhibitors were prepared as a 20 mM stock solution and diluted to working concentrations in the same BLI binding buffer by serial dilution. After establishing a baseline with the binding buffer alone, biosensors were immersed in wells containing varying concentrations of MYCi during the association step and subsequently transferred to wells with binding buffer alone during the dissociation step. All raw BLI binding data were collected and processed using ForteBio Data Analysis software (v7.0).

### Pull-down assay

HA-MYC-WT (1-439) and HA-MYC deletion mutants (ΔA: 1-63, ΔB: 64-126, ΔC: 127-189 [eMBII], ΔD: 190-252, ΔE: 253-315, ΔF: 316-378, ΔG: 379-439 [bHLH-LZ]) were kind gifts from Prof. William P. Tansey (Vanderbilt University School of Medicine, Nashville, TN) (Table S1). Plasmids were transfected with Lipofectamine 2000 (Thermo Fisher Scientific, 11668-019) in HEK293 cells, according to the manufacturer’s instructions. After 48-hour transfection, cell pellets were collected. Collected HEK293 cell pellet (2 x10^6^ cells per pulldown condition) was suspended in 300 µL pulldown lysis buffer (50mM Tris-HCl, pH 7.4, 150 mM NaCl, 1mM EDTA, 0.1% IGEPAL + 1x protease/phosphatase inhibitor) and lysed by repeat (3x) freeze-thawing in liquid nitrogen. The lysates were spun at 4°C for 15 minutes at 15,000 rpm. The collected supernatants were pre-cleared with Pierce streptavidin magnetic beads (Thermo Fisher Scientific, 88817) for 1 hr at 4°C. Approximately 300 µg protein was applied to each sample and incubated with 10 µM of biotinylated-MYCi975 or D-biotin on a rotator overnight at 4°C. Next day, 60 µL of streptavidin beads were added to each sample and further rotated for 1 hour at 4°C. Beads were washed with wash buffer (lysis buffer containing 0.1% BSA) for 3 times and another 3 times with lysis buffer, then eluted with 50 µL of 2x Laemmli sample buffer and boiled at 95°C for 5 min. The supernatant was subjected to western blot.

### Protein expression and purification

Full-length human c-MYC and MYC variants (Table S2) were cloned into a modified pET-28a (+) vector (GenScript) containing an N-terminal hexahistidine (His) tag followed by a TEV protease cleavage site (ENLYFQG). All constructs were expressed in Escherichia coli BL21-Gold (DE3) (Agilent, 230132). DE3 cells (100 µL) were transformed with 100 ng plasmid via heat shock (42°C, 30 sec) after incubation on ice (30 min). Cells were recovered in 0.9 mL LB medium (1 hr, 37°C, 225–250 rpm), pelleted (1,000 × g, 4 min), and plated on LB agar supplemented with 50 µg/mL kanamycin (overnight, 37°C). Single colonies were inoculated into 5 mL LB-kanamycin (50 µg/mL) and grown overnight (37°C, 200 rpm). Cultures were scaled up to 30 mL (6× dilution) in fresh LB-kanamycin and grown overnight. A final 10× expansion (300 mL total) in antibiotic-free LB was grown to mid-log phase (OD_600_ = 0.6–0.8), and protein expression was induced with 0.5 mM IPTG (6 hr, 37°C). Cell pellets were resuspended in denaturing lysis buffer (8 M urea, 100 mM NaH□PO□, 10 mM Tris, pH 8.0; 12 mL per 300 mL culture) and incubated for 1 hr (room temperature, 260 rpm). Cleared lysates were obtained by centrifugation (10,000 ×g, 30 min, RT) and incubated with Ni-NTA agarose slurry (Qiagen, 30210; 1 mL slurry per 4 mL lysate) for 1 hr (RT, 260 rpm). His-tagged proteins were purified using 5 mL polypropylene columns (Qiagen, 34964) with imidazole gradient elution. Proteins were desalted into BLI assay buffer (50 mM HEPES, 150 mM NaCl, pH 7.4) using Zeba Spin columns (Thermo Fisher, 89892) following the manufacturer’s protocol.

### NMR spectroscopy

Uniformly ^15^N/^13^C-labeled bHLH-LZ domain protein samples were prepared as previously reported by Panova et al., 2019 (*31*). Briefly, the protein was expressed in E. coli BL21(DE3) cells. Cell pellets were resuspended in lysis buffer containing 20 mM sodium phosphate, 8 M urea, pH 7.4. Cells were lysed by sonication while stirring on ice using a 5s on/10 s off cycle at 35% amplitude for a total of 5 minutes, repeated twice. The lysate was clarified by centrifugation at 24,424 x g for 60 minutes, and the supernatant was filtered. The supernatant was then loaded onto a pre-equilibrated HiTrap TALON crude column (Cytiva) using the Cytiva AKTA GO system. The flow rate was kept slow to maximize protein-resin interaction. The column was first washed with buffer containing 3 M guanidine-HCl, 20 mM sodium phosphate, 0.1 M NaCl, pH 7.4, followed by a secondary wash using a buffer containing 2.5 mM imidazole, 20 mM sodium phosphate, 0.1 M NaCl, pH 7.4. The protein was eluted with an elution buffer containing 125 mM imidazole, 20 mM sodium phosphate, 0.1 M NaCl, pH 7.4. The elution profile was monitored by the AKTA system, and protein-containing fractions were collected based on absorbance at 280 nm. The pooled fractions were concentrated and subjected to size-exclusion chromatography (SEC) on a Superdex 200 (S200) column, equilibrated with 20 mM sodium phosphate (pH 7.4), 150 mM NaCl, 1 mM DTT, and 0.05% (w/v) sodium azide.

### Sample Preparation and NMR Data Acquisition

Purified protein was concentrated to 200 µM in the final SEC buffer supplemented with 10% (v/v) D_2_O. For interaction studies, MYCi inhibitors were dissolved in d6-DMSO and added to the protein samples to a final molar ratio of 1:2 (protein: ligand). The eMBII peptide was prepared in the same buffer and titrated into the bHLH sample at the same 1:2 molar ratio. 2D ^1^H-^15^N HSQC NMR experiments were performed at 298 K on a Bruker 600 MHz Avance Neo spectrometer, equipped with a cryogenic quadruple-resonance probe, located at the Northwestern University IMSERC facility. All NMR data were processed using NMRPipe and analyzed using POKY software (*32, 33*).

### Molecular Docking

AlphaFold 3 was used to individually generate models of MYC bHLH-LZ domain (amino acids 353-439) and the eMBII domain (amino acids 127-189) (*23*). Unrestrained docking calculations between bHLH-LZ, eMBII, and MYCi975 were carried out using the HADDOCK (High Ambiguity Driven protein-protein DOCKing) server (v2.4) (*34, 35*). All parameters were set to default values except the number of structures of rigid body docking (value set to 1000), number of structures for final refinement (value set to 400) and minimum cluster size to 1. A total of 400 models were generated for each complex clustered using Fraction of Common Contacts (FCC) of 0.6. The top scoring model/clusters were then visually assessed for compatibility with i) key residues identified from BLI, and ii) previously published structure-activity relationship studies indicating that the chlorobenzyl ring is not required for high affinity complexation (*10*).

### CRISPR tiling screen

A CRISPR tiling screen targeting the human MYC gene was conducted using 1000 sgRNA sequences (222 sgRNAs saturating the entire MYC coding sequence, the rest of guides targeting introns and 500 bp putative MYC regulatory region upstream of first exon for a total of 886 guides; as well as 114 nontargeting control guides), designed via the DESKGEN platform (Desktop Genetics, Inc.). PC3M cells were first transduced with MSCV-hCas9-PGK-Puro (Addgene, #65655) and selected with puromycin to generate the PC3M-Cas9+ cell line used for library transduction. The guides were selected based on specificity (Hsu 2013 specificity score ≥50) (*36*) and predicted activity (Doench 2016 activity score) (*37*), to ensure on-target efficiency. sgRNA oligonucleotides were synthesized and cloned into the LRG (Lenti_sgRNA_EFS_GFP, Addgene, 65656) lentiviral vector, via digestion with BsmBI and insertion of oligo duplex via Gibson assembly. Lentiviral particles were produced by transfecting HEK293 cells with psPAX2 (Addgene, 12260) and pMD2.G (Addgene, 12259), followed by titration to achieve a 0.3 multiplicity of infection (MOI), confirmed by flow cytometry for GFP expression. To ensure a minimum of 10,000× cells/sgRNA representation, 6 × 10□ PC3M-Cas9+ cells were transduced with the MYC tiling library providing approximately 60,000× initial coverage. Transduced cells were selected with puromycin (2 µg/mL; Gibco, A1113803) for 3 days, followed by a 9-10-day selection/cutting period. Throughout the screen, cells were passaged every 2-3 days and maintained at a representation of ≥10,000 cells per sgRNA. Afterward, cells were split and exposed to either 6 µM MYCi975 or DMSO for 35 days, with fresh drug replenished every 3-4 days, to assess the effects of MYC perturbation on guide selection. Nontargeting control guides were included to account for baseline proliferation and toxicity effects. The screen was performed in two biological replicates by two different investigators. Collected cell pellets on Day 21 and Day 35 were submitted for sequencing.

On the 21^st^ and 35^th^ days of post-transduction, raw sequencing data were collected and analyzed to evaluate the relative abundance of each sgRNA targeting different regions of the *c-MYC* protein-coding sequence in both untreated and MYCi975-treated (6µM) cohorts. Genomic DNA extraction was performed using Zymo DNA Miniprep Plus kit (Zymo research, D4086) and used to amplify integrated sgRNA sequences using library prep method described previously for LentiGuide-Puro vector (*38*). The barcoded sequencing libraries were then separated on agarose gel and the 350bp target amplicon purified using NucleoSpin Gel and PCR Clean-up Mini kit (Macherey-Nagel, 740609.50). Libraries were pooled and sequenced on Illumina NovaSeq X Plus. The sgRNA sequences were aligned to the reference *c-MYC* functional region, and for each sgRNA at both time points, the log10 fold change in normalized abundance between MYCi975-treated and control conditions was calculated. This data was further standardized by computing Z-scores. The results were then visualized using R to pinpoint key protein regions critical for cellular proliferation and survival in the presence of MYCi975, highlighting the most functionally significant segments of the *c-MYC* gene under MYCi975-treated conditions. The entire screen was carried out independently by two distinct investigators.

### Validation of CRISPR tiling screen

To validate the CRISPR tiling screen of MYC, top-ranked sgRNAs targeting the MYC eMBII and bHLH domains were synthesized and cloned into the lentiGuide-Puro vector (Addgene, 52963) (Table S3). Plasmids were amplified and purified using the Maxiprep Kit (Thermo Fisher Scientific, K210007). PC3M-Cas9-expressing cells were co-transfected with sgRNA plasmid pairs (12 µg plasmid) using Lipofectamine 3000 (Invitrogen, L3000015) in Opti-MEM medium (Thermo Fisher Scientific, 31985062). Transfected cells underwent selection with puromycin 2 µg/mL (Gibco, A1113803) for three days and were cultured for 9–10 days to allow for genomic editing. Subsequently, cells were treated with MYCi975 to identify mutations conferring resistance. The MYC locus was sequenced to delineate resistance-associated mutations within the eMBII and bHLH domains.

### Targeted ultra-deep sequencing

Genomic DNA was extracted from PC3M-Cas9 cell populations (sgRNA library-transfected and selected with MYCi975 for 35 days, or co-transfected with specified guide pairs A169+A361 and A169+A423) using the Zymo DNA Miniprep Plus kit (Zymo Research, D4086). Total RNA was isolated in parallel from the same samples using the RNeasy Mini Kit (Qiagen, 74134), including on-column DNase I digestion. Nucleic acid concentration and purity were assessed by Qubit fluorometry. For RNA samples, 1 µg of total RNA was reverse transcribed into cDNA using the iScript cDNA Synthesis Kit (Bio-Rad, 1708891) according to the manufacturer’s protocol. For each sample, 200 ng of cDNA of MYC encompassing the eMBII and bHLH-LZ regions were enriched by a custom two-step PCR approach. First, target regions were amplified from 50 ng of prepped libraries using a primer pool (Integrated DNA Technologies) with KAPA HiFi HotStart ReadyMix (Roche, KK2602). A second, 8-cycle indexing PCR was performed to attach full Illumina sequencing adapters. The Qiagen QIAseq 1-step Amplicon Library kit (Qiagen 180412) and the Qiagen Y-UDI Adapters (Qiagen, 180310) were used to add index barcode sequences to the libraries. Libraries were quantified by qPCR (KAPA Library Quantification Kit), normalized, pooled equimolarly, and sequenced on an Illumina MiSeq platform (MiSeq Reagent Kit v3, 600-cycle) to generate 2×300 bp paired-end reads, achieving >10,000× average depth across the targeted MYC locus.

Raw sequencing reads were demultiplexed using Illumina bcl2fastq (v2.20) with default parameters. Adapter sequences and low-quality bases (Q < 20) were trimmed using Trim Galore (v0.4.4). Cleaned reads were aligned to the human reference genome (GRCh38/hg38) using BWA-MEM (v0.7.12). Duplicate reads were marked with Picard Tools (v2.17.6). Local realignment around indels and base quality score recalibration was performed using GATK (v4.1.0). Variants (single-nucleotide variants and indels) were called across the targeted MYC region using bcftools (v1.10.1) mpileup with parameters -B-q 20-Q 20-d 100000, followed by BCFtools call with the multiallelic-caller model. Variants were filtered to retain those with a minimum read depth of 100×, allele frequency ≥0.5%, and Fisher’s strand bias p-value > 0.01. Variant annotation was performed using SnpEff (v4.3k) with the hg38 database.

### Generation of stable cell lines

MYC and MYC mutants containing point mutations near eMBII (SVF-WAL) and in bHLH (RSF-QRA) domains were synthesized and cloned into the pCDH-MCS-T2A-Puro-MSCV vector (GenScript). Plasmids were amplified and purified using a Maxiprep kit (Thermo Fisher Scientific, K210007). Lentiviral particles were produced in HEK293 cells using a third-generation packaging system. The packaging plasmids pMDLg/pRRE (Addgene, 12251), pRSV-Rev (Addgene, 12253), and pMD2.G (Addgene, 12259) were co-transfected with 12 µg of MYC, WAL MYC, QRA MYC and WAL-QRA MYC mutant plasmids using Lipofectamine 3000 (Invitrogen, L3000015) in Opti-MEM medium (Thermo Fisher Scientific, 31985062). Viral supernatant was harvested 48 hours post-transfection, 22Rv1, PC3 and HO15.19 MYC-null fibroblast cells were then transduced with the harvested lentiviral particles to achieve stable integration of the plasmids. At 48 hours post-transduction, cells were selected with 2 µg/mL puromycin (Gibco, A1113803) for 7–10 days to establish stable cell lines. These stable lines were subsequently used for treatment with the MYCi975.

### Clonogenic survival assay

To assess long-term proliferative capacity and MYCi sensitivity, clonogenic assays were performed. Cells were seeded at low density (typically 300 cells per well) in six-well plates. Following cell attachment, the medium was replaced with a fresh medium containing either vehicle control (DMSO) or varying doses of MYCi975 and MYCi648. Cells were incubated for 14 days to allow for colony formation, with the medium (including treatment) replenished every 3 to 4 days. After the incubation period, colonies were fixed with 4% paraformaldehyde or methanol for 15 minutes and then stained with a 0.5% crystal violet solution for 30 minutes. Excess stain was removed by thorough washing with distilled water, and plates were air-dried. Colonies were imaged using a standard scanner or digital camera. Quantification was performed by analyzing the stained images with ImageJ software.

### Fluorescence in situ hybridization (FISH)

FISH analysis was performed on PC3M cultured cells, which were harvested following standard laboratory procedures. The MYC dual-color break-apart probe (Abbott Park, Illinois) labeled the 5’MYC as spectrum red and the 3’MYC as spectrum green. The normal signal pattern shows two fusions (overlapping green and red signals). Single green and/or red signals indicate MYC rearrangements, while three or more fusions indicate an increased copy number. To establish the threshold for true amplification, baseline MYC copy numbers were analyzed in 20 normal control samples. Based on this analysis where 0% of normal cells exhibited ≥ 5% copies, a cut-off of five or more MYC copies per cell was defined as the criterion for MYC amplification. A total of 100 cells were counted independently by two technologists under a fluorescent microscope, and images were captured at 100x magnification following standard procedures.

### Acetylated-K148/K157-MYC expression and purification

Acetylated MYC (K148/K157) was expressed using a two-plasmid system containing pRSFDuet-1/AcKRS3/PylT and pTech/AcKRS/PylTopt. These plasmids, generously provided by Dr. Michael Lammers from the University of Greifswald, facilitate the site-specific incorporation of acetylated lysine residues during protein expression. The MYC expression constructs, containing K148/K157 substitutions, were cloned into pET30a vectors with an N-terminal hexa-histidine tag followed by a TEV protease cleavage site. Protein expression was performed in BL21(DE3)-CodonPlus cells following established protocols (*24*). After expression and purification of acetylated K148/K157 MYC, the expressed proteins were filtered through Amicon 30kDa Centrifugal Filter (Millipore, UFC8030) to remove truncated forms of the expressed MYC.

Site-specific incorporation of acetylated K148/K157 in recombinant MYC was confirmed via LC-MS/MS. Peptides were separated using an increasing gradient of organic mobile phase B (80 % acetonitrile, 0.1 % aqueous formic acid) on a Thermo Fisher Vanquish Neo nano UHPLC system. A linear 45-minute gradient from 1 % to 40% (v/v) B eluted peptides from a Thermo Scientific Acclaim Pepmap column (75 μm × 15 cm, C18) at 350 nl/min flow rate. A Thermo Scientific Orbitrap Exploris 480 mass spectrometer acquired full scan MS1 data (120k resolution, 350-1600 m/z mass range, automatic maximum injection time) and MS2 data (15k resolution, 1.4 m/z isolation width, 30 % HCD collision energy, automatic maximum injection time) to produce raw files. Raw files from MYC acetylated and control samples were processed using FragPipe (v22.0) using the PTM search workflow. Data was searched against the UniProt E. coli BL21 (DE3) database, supplemented with the MYC sequence (P01106-1). The following modifications were considered as variable: acetylation of lysine and the N-terminus (mass shift of 42.0106 Da), and oxidation of methionine (mass shift of 15.9949 Da), with a maximum of three variable modifications. PTMProphet and MSBooster were enabled, with the Koina server (https://koina.wilhelmlab.org:443/v2/models/) utilized for peptide scoring. The resulting output file, “combined_modified_peptide.tsv,” was processed using a custom R script to filter for peptides mapped to P01106-1 with acetylation. Specific peptide sequence patterns (LVSEK, LASYQAARK, and KDSGSPNPA), which correspond to regions containing the target modification sites were retained. Precursor ion intensities for these peptides were visualized using Skyline software.

### AlphaFold 3 modeling of unacetylated and acetylated MYC

From the AlphaFold 3 server, the top five predicted models were downloaded for unacetylated MYC and MYC with N^ε^-acetylation of lysine at residues 148 and 157. For each model, Predicted Local Distance Difference Test (pLDDT) scores were averaged per amino acid to generate mean pLDDT scores across the 439-amino acid protein, which were subsequently plotted for each model. Additionally, Predicted Aligned Error (PAE) values for each amino acid pair were extracted.

### Immunoprecipitation and western blot analyses

22Rv1 cells were lysed using a lysis buffer (20 mM Tris-HCl [pH 8.0], 100 mM NaCl, 1 mM EDTA, 0.5% Nonidet P-40). The insoluble fraction was treated with an enzymatic shearing cocktail from the Nuclear Complex Co-IP Kit (Active Motif, 54001) at 4°C for 90 minutes to release nuclear proteins. Soluble and insoluble fractions were combined, and 500 µg of total protein was incubated overnight at 4°C with 10 µM Biotin MYCi975, 10 µM D-biotin following the Dynabeads® Co-Immunoprecipitation Kit protocol (Thermo Fisher Scientific, 14321D). After incubation, samples were washed three times with lysis buffer, once with wash buffer, and eluted with the provided elution buffer. Eluted proteins were analyzed by western blotting. Western blot analysis was performed as previously described (*10, 39*). Primary antibodies used including MYC (Y69; Abcam, ab32072), HA-tag (C29F4) (Cell Signaling, 3724S), c-Myc (9E10) (Santa Cruz, SC-40), Anti-acetyl-c-Myc (Lys148) Antibody (Abe25, Millipore), β-actin (Cell Signaling, 5125S), GAPDH (Cell Signaling, 3683S) and Anti-FLAG® (Sigma-Aldrich, F1804).

### MYC acetylation in human samples

To investigate the post-translational modification (PTM) landscape of the MYC protein specifically focusing on K148 lysine acetylation, high-resolution acetylome data were retrieved from the National Cancer Institute (NCI) Proteomic Data Commons (PDC) (https://pdc.cancer.gov/pdc). The analysis incorporated diverse cancer cohorts from the Clinical Proteomic Tumor Analysis Consortium (CPTAC) to capture the heterogeneity of MYC regulation across various malignancies. Specifically, the following cohorts were integrated for detailed analysis: Lung Adenocarcinoma (PDC000491 and PDC000224) (*40, 41*), Lung Squamous Cell Carcinoma (PDC000233) (*42*), Breast Invasive Carcinoma (PDC000239) (*43*), Glioblastoma Multiforme (PDC000450, PDC000245) (*44, 45*), Uterine Corpus Endometrial Carcinoma (PDC000443 and PDC000226) (*46, 47*) and quantitative acetyl site-level data were processed based on the MYC protein sequence (Accession: ENSP00000259523.6 and NP_001341799.1). Our analytical focus was directed toward the K148 residue in isoform 1 (P01106-1), located within the conserved peptide motif LVSEKLASYQAAR. This site was identified across multiple patient biospecimens as a critical regulatory node. To ensure data comparability across different Tandem Mass Tag (TMT) plexes and experimental batches, acetylation intensities were normalized and expressed as log_2_ TMT ratios. These values were used to perform comparative assessments between different tumor tissues and matched normal tissues, facilitating the identification of tumor-specific accumulation of K148-MYC acetylation patterns.

### Cell viability

Cells were seeded in 96-well plates at a density of 3,000 cells per well and incubated for 24 hours to allow attachment. For endpoint cell viability assessment, 22Rv1, PC3, PC12, MV-4:11, HL-60, THP-1, Kasumi-1, HT-29, HepG2, MDA-MB-231, A549, A2780, and PLC/PRF/5 cells were treated in quadruplicate with serial dilutions of MYCi975 or DMSO vehicle control. After 72 hours, cell viability was determined using the CellTiter-Glo Luminescent Assay (Promega, G7573), with luminescence signals normalized to vehicle controls for IC calculation via nonlinear regression. In parallel, real-time viability kinetics were assessed in HO15.19 MYC and HO15.19 WAL-QRA MYC cells following treatment with varying MYCi975 concentrations. These cells were monitored every 24 hours for 96 hours using the IncuCyte S3 live-cell analysis system, and viability was determined by quantifying total cells normalized to the 0-hour tim point.

### Synthetic procedures for MYCi648

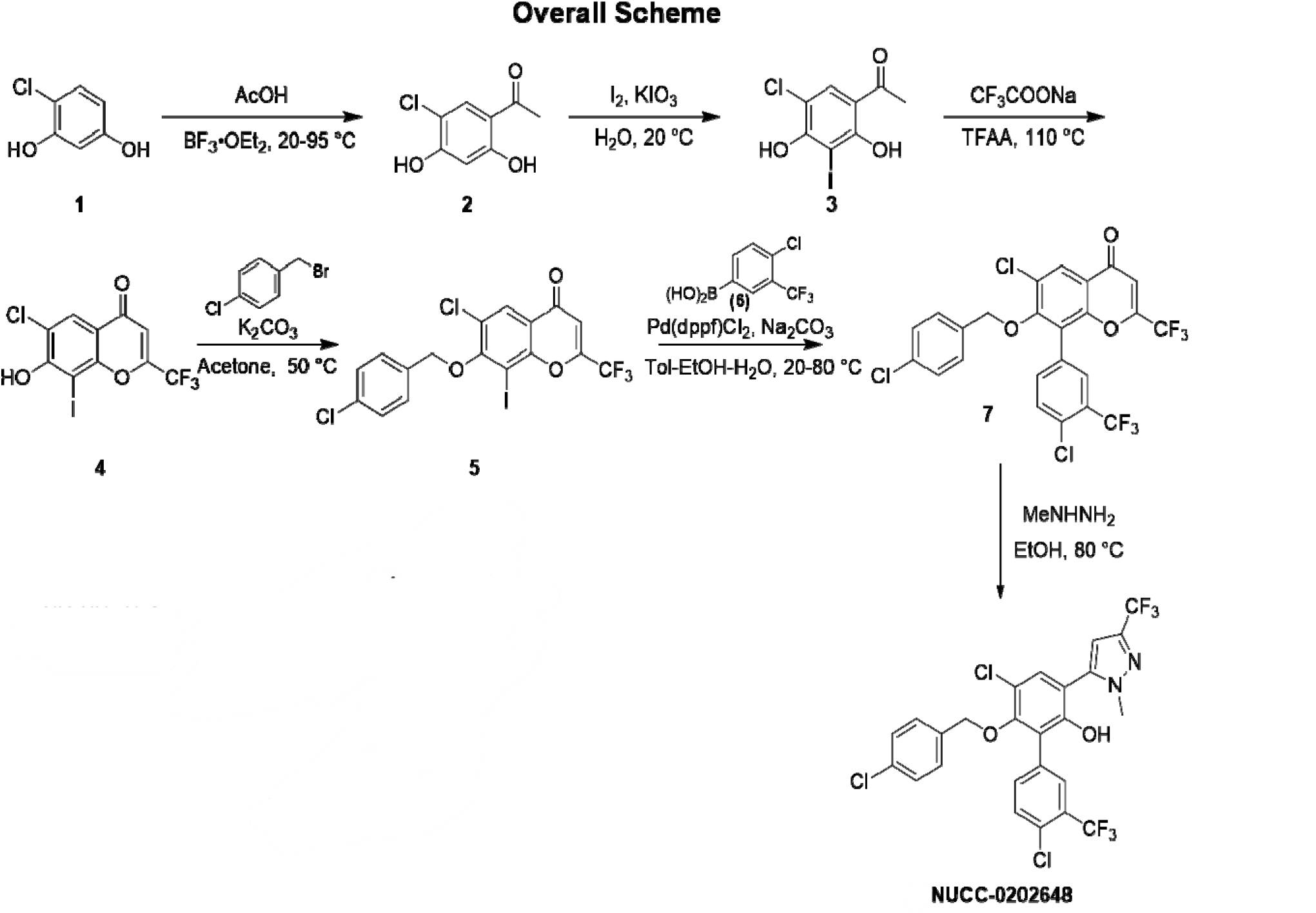

### Synthesis of 1-(5-chloro-2,4-dihydroxyphenyl) ethan-1-one (2)

To a suspension of 4-chlorobenzene-1,3-diol (1, 10 g, 69.18 mmol) in BF3•Et2O (50 mL) wa added AcOH (4.15 g, 69.18 mmol) drop-wise under N2 at 20 °C. The mixture was heated to 95 °C for 5 hours. A large quantity of yellow solid was formed. The mixture was cooled to 20 °C and poured into a solution of AcONa (10%, 160 mL). The mixture was stirred at 20 °C for 3 hours to produce copious solids. LC-MS showed that 71% of desired product mass was detected. The solid was collected by filtration. The solid was dissolved in ethyl acetate (100 mL) and dried with anhydrous Na2SO4, filtered, and concentrated. The residue was purified by prep-HPLC (formic acid modifier) to afford compound 2 (7.9 g, 60% yield) as a yellow solid. 1H NMR (400 MHz, DMSO-d6) δ: 12.34 (s, 1H), 11.40 (s, 1H), 7.87 (s, 1H), 6.46 (s, 1H), 2.54 (s, 3H). ESI-MS: m/z = 187.2 ([M + H]+).

### Synthesis of 1-(5-chloro-2,4-dihydroxy-3-iodophenyl) ethan-1-one (3)

To a solution of compound 2 (6.9 g, 36.61 mmol) in EtOH (60 mL) and H2O (96 mL) was added I2 (4.51 g, 17.77 mmol) and potassium iodate (1.58 g, 7.38 mmol) at 20 °C. The mixture wa stirred at 20 °C for 16 hours after which copious yellow solids formed. LC-MS showed compound 2 was consumed completely, and 93% of desired compound mass was detected. The solid was collected by filtration. The solid was dissolved in ethyl acetate (100 mL) and washed with brine (50 mL). The ethyl acetate phase was separated and dried with dried with anhydrous Na2SO4, filtered, and concentrated to afford compound 3 (9.95 g, 88% yield) as a brown solid. 1H NMR (400 MHz, DMSO-d6) δ: 13.51 (s, 1H), 11.24 (s, 1H), 8.03 (s, 1H), 2.62 (s, 3H). ESI-MS: m/z = 313.2 ([M + H]+).

### Synthesis of 6-chloro-7-hydroxy-8-iodo-2-(trifluoromethyl)-4H-chromen-4-one (4)

To a solution of compound 3 (2.4 g, 7.60 mmol) in TFAA (12 mL) was added sodium 2,2,2-trifluoroacetate (2.28 g, 16.73 mmol) at 20 °C. The mixture was stirred at 110 °C for 60 hours. LC-MS showed compound 3 was consumed completely, and 68% of desired compound mass was detected. The reaction mixture was cooled to 20 °C and poured into water (200 mL). The mixture was extracted with ethyl acetate (200 mL×2). The combined organic layers were washed with brine (200 mL), dried over Na2SO4, filtered, and concentrated. The residue was purified by column chromatography (SiO2, petroleum ether/ethyl acetate = 10:1 to 5:1) to afford compound 4 (2.1 g, 69% yield) as a white solid. 1H NMR (400 MHz, DMSO-d6) δ: 7.99 (s, 1H), 7.03 (s, 1H). ESI-MS: m/z = 391.0 ([M + H]+).

### Synthesis of 6-chloro-7-((4-chlorobenzyl) oxy)-8-iodo-2-(trifluoromethyl)-4H-chromen-4-one (5)

To a solution of compound 4 (1 g, 2.54 mmol) and 1-(bromomethyl)-4-chloro-benzene (781.44 mg, 3.80 mmol) in acetone (10 mL) was added K2CO3 (1.05 g, 7.61 mmol) at 20 °C. The mixture was heated to 50 °C and stirred for 14 hours. LC-MS showed compound 4 was consumed completely. The reaction mixture was cooled to 20 °C and poured into water (50 mL). The mixture was extracted with ethyl acetate (50 mL×2). The combined organic layers were washed with brine (50 mL), dried over Na2SO4, filtered, and concentrated. The residue was purified by column chromatography (SiO2, petroleum ether/ethyl acetate = 80:1 to 50:1) to afford compound 5 (1 g, 65% yield) as a yellow solid. 1H NMR (400 MHz, DMSO-d6) δ: 8.14 (s, 1H), 7.65-7.62 (m, 2H), 7.54-7.51 (m, 2H), 7.15 (s, 1H), 5.12 (s, 2H). ESI-MS: m/z = 514.9 ([M + H]+).

### Synthesis of 6-chloro-8-(4-chloro-3-(trifluoromethyl) phenyl)-7-((4-chlorobenzyl)oxy)-2-(trifluoromethyl)-4H-chromen-4-one (7)

A mixture of compound 5 (500 mg, 834.87 umol), (4-chloro-3-(trifluoromethyl)phenyl)boronic acid (6, 206.05 mg, 918.36 µmol), Na2CO3 (176.97 mg, 1.67 mmol) and Pd(dppf)Cl2 (61.09 mg, 83.49 umol) in toluene (6 mL), EtOH (1 mL), and H2O (2 mL) was degassed and purged with N2 for 3 times at 20 °C. The reaction mixture was heated to 80 °C and stirred for 16 hours under N2 atmosphere. LC-MS showed compound 5 was consumed completely. The reaction mixture was cooled to 20 °C and poured into water (50 mL). The mixture was extracted with ethyl acetate (50 mL×2). The combined organic layers were washed with brine (50 mL), dried over Na2SO4, filtered, and concentrated. The residue was purified by column chromatography (SiO2, ethyl acetate), then purified by prep-HPLC (TFA modifier) to afford compound 7 (290 mg, 59% yield) as an off-white solid. ESI-MS: m/z = 567.1 ([M + H]+).

### Synthesis of 4’,5-dichloro-6-((4-chlorobenzyl) oxy)-3-(1-methyl-3-(trifluoromethyl)-1H-pyrazol-5-yl)-3’-(trifluoromethyl)-[1,1’-biphenyl]-2-ol (9, MYCi648)

To a solution of compound 7 (220 mg, 379.78 µmol) in EtOH (2 mL) was added MeNHNH2 (131.23 mg, 1.14 mmol). The mixture was stirred at 80 °C for 1 hour. LC-MS showed compound 7 was consumed completely, and two peaks with desired product mass (40% and 57%) were detected. The reaction mixture was concentrated. The residue was purified by prep-HPLC (TFA modifier) to afford compound 9, MYCi648 (48.2 mg, 20% yield) as a brown solid and the pyrazole methyl isomer (60.3 mg, 26% yield) as a white solid. 1H NMR (400 MHz, DMSO-d6) δ: 9.44 (s, 1H), 7.76-7.73 (m, 2H), 7.65 (dd, J = 8.4, 2.0 Hz, 1H), 7.55 (s, 1H), 7.29-7.26 (m, 2H), 6.99-6.96 (m, 2H), 6.84 (s, 1H), 4.69 (s, 2H), 3.78 (s, 3H). ESI-MS: m/z = 594.9 ([M + H]+).

### Pharmacokinetic Studies

Pharmacokinetic studies on MYCi648 and MYCi975 were performed by Sanford Burnham Prebys Medical Discovery Institute, with mice strains C57BL/6 male and CD-1 male, respectively. MYCi648 was formulated in PBS with 10% DMSO and 20% Tween-80 and 50 mg/kg was given at a single dose either via i.p. or p.o. routes. MYCi975 was formulated in corn oil with 5% DMSO for i.p. and p.o. dosing (100 mg/kg and 250 mg/kg), and in 10% HPBCD with 5% ethanol and 2% Tween80 for i.v. dosing (2 mg/kg). Each data point represents the mean value from three experimental mice.

### Statistical Analysis

Descriptive statistics were presented as mean ± SD or mean ± SEM. The normality of data distribution was assessed using the Shapiro-Wilk test. For continuous data, two-tailed Student’s t-test was used when normally distributed, while two-way ANOVA followed by Tukey’s post hoc test was applied for multiple comparisons. Non-normally distributed data were analyzed using the Mann-Whitney U test. All statistical analyses were two-tailed, with a significance level of p<0.05 (represented as *p<0.05, **p<0.01, ***p<0.001, ****p<0.0001). All statistical analyses were performed using licensed GraphPad Prism (v9.2).

## Acknowledgments

We thank Northwestern NUSeq, Keck Facility Core of the Northwestern Center, and the Northwestern Proteomics Core Facility, generously supported by NCI CCSG P30 CA060553 awarded to the Robert H Lurie Comprehensive Cancer Center, instrumentation award (S10OD025194) from NIH Office of the Director, and the National Resource for Translational and Developmental Proteomics supported by P41 GM108569. We further acknowledge the support of Northwestern NUCures program (NU2015-150).

## Funding

This work was supported by:

National Institutes of Health grant P50 CA180995 (S.A.A.).

National Institutes of Health grant R01 CA257258 (S.A.A., D.C.).

National Institutes of Health grant R35 GM143054 (J.J.Z).

Prostate Cancer Foundation (PCF) TACTICAL Award (S.A.A.).

Polsky Urologic Cancer Institute (S.A.A.).

## Author contributions

**Conceptualization:** D.G.G., M.I.T., and S.A.A.

**Methodology:** D.G.G., M.I.T., and S.A.A.

**Investigation:** D.G.G., M.I.T., A.W.T.S., J.B.P., D.H.R., S.Q., A.A.E., H.P., W.Y., X.L., K.U., M.M.K., C.E.B., M.F.D., K. K., J.S., M.L., G.E.S., J.J.Z., and S.A.A.

**Visualization:** D.G.G., M.I.T., and S.A.A.

**Funding acquisition:** S.A.A., D.C., and J.J.Z.

**Project administration:** M.M.K., and S.A.A.

**Supervision:** D.C., G.E.S., J.J.Z., and S.A.A.

**Writing – original draft:** D.G.G., and S.A.A.

**Writing – review & editing:** All authors.

## Competing interests

S.A.A. and G.E.S. are co-founders of Vortex Therapeutics, which is licensing Intellectual Property (IP) related to the science and materials found in this manuscript. S.A.A., G.E.S, H.H., and M.I.T. are co-inventors on that same IP. All other authors declare that they have no competing interests.

## Data and materials availability

The proteomic and acetylproteomic datasets analyzed in this study are publicly available through the Proteomic Data Commons (PDC) (https://pdc.cancer.gov/pdc). The specific data subsets can be accessed using the following PDC Study IDs: PDC000491, PDC000224, PDC000443, PDC000233, PDC000226, PDC000245, and PDC000239.

---

**Figure S1.**
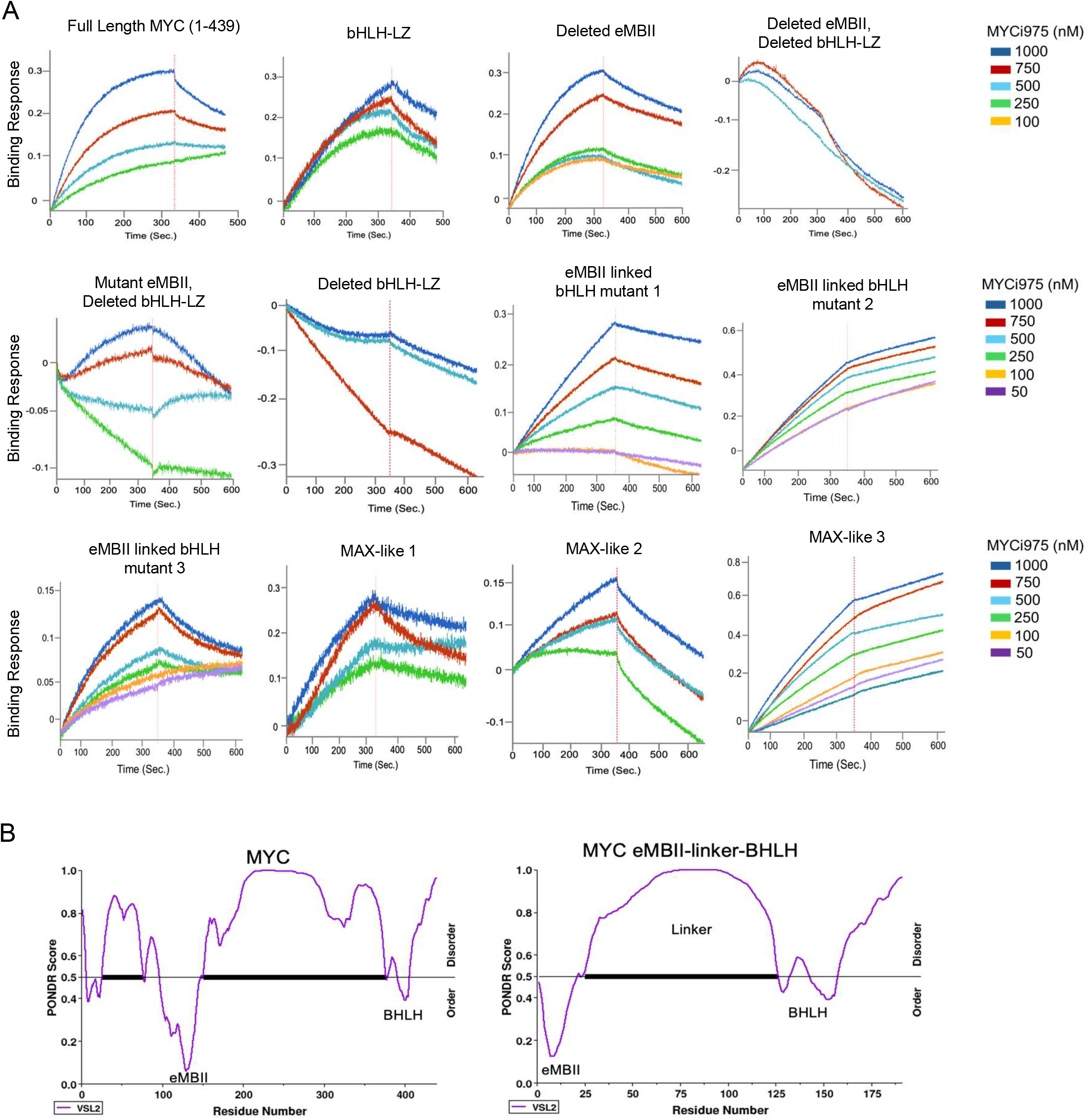
**(A)** BLI sensorgrams of MYCi975 binding to WT and mutant MYC proteins. **(B)** PONDR-VSL2 disorder predictions of full-length MYC and the MYC eMBII linked to bHLH construct.

**Figure S2.**
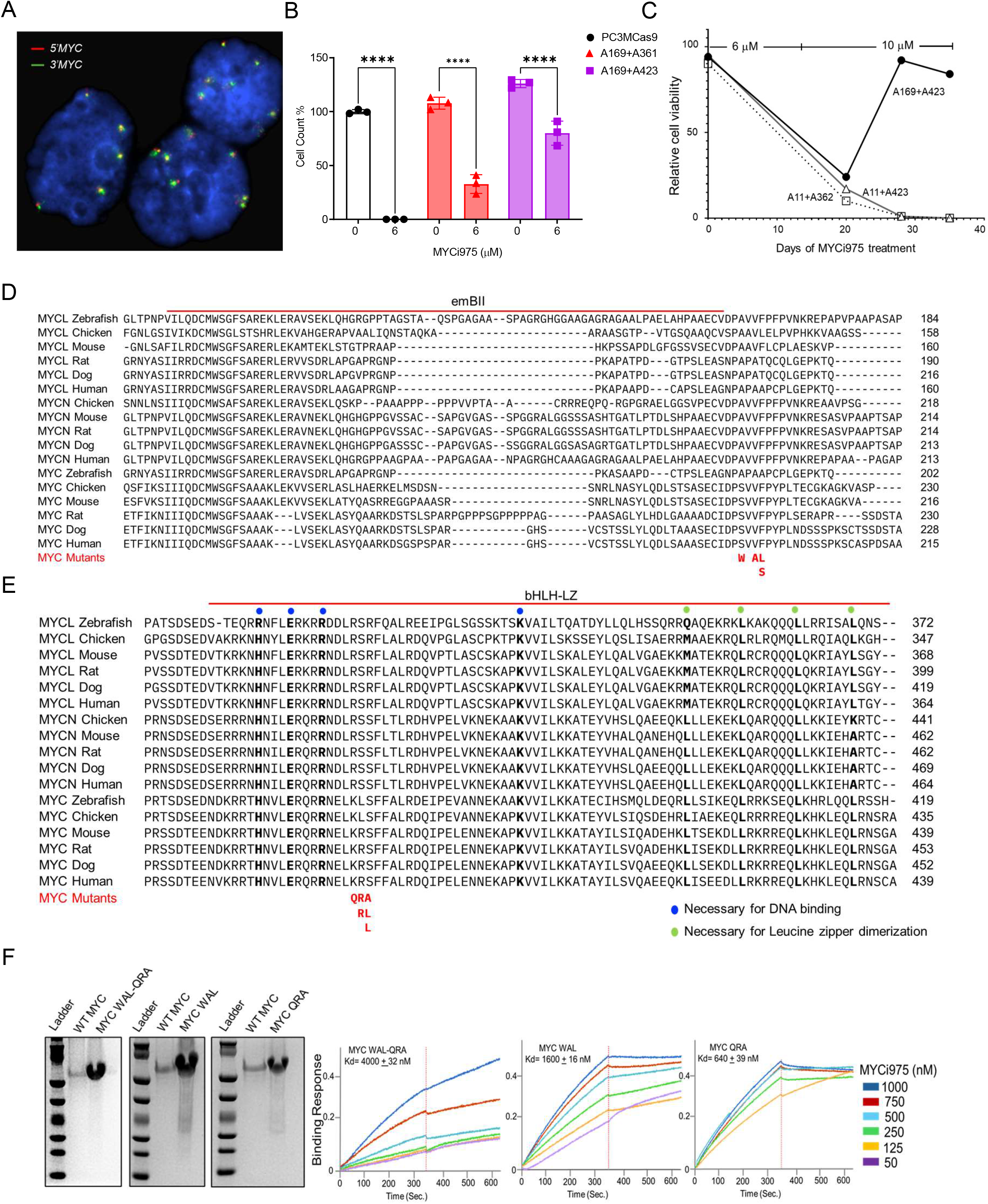
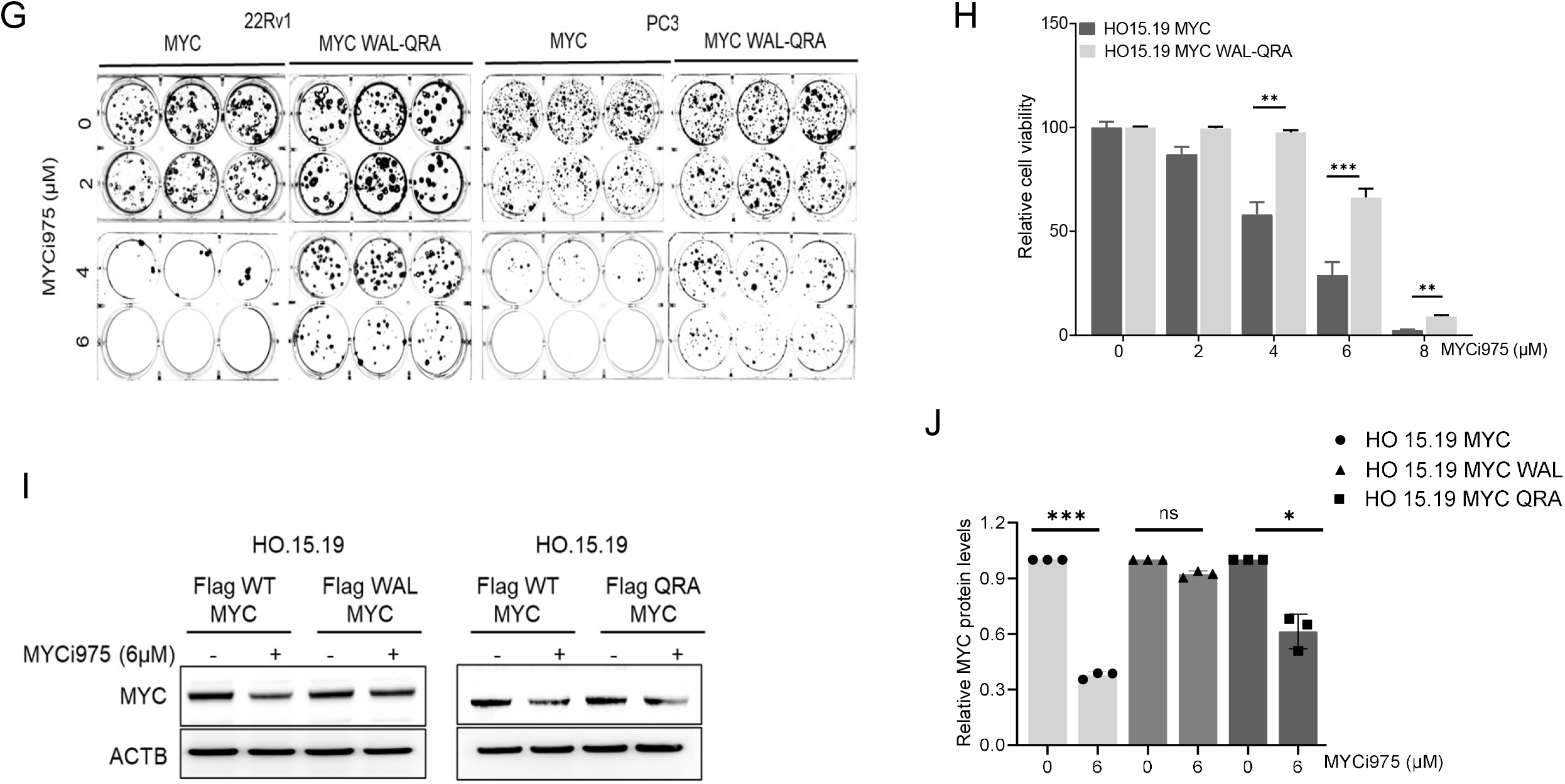
Functional and evolutionary analyses of MYC residues required for MYCi975 binding. **(A)** Representative FISH image of PC3M cells. The MYC dual-color break-apart probe (Abbott Park, Illinois) labeled the 5’MYC as red and the 3’MYC as green. The normal signal pattern shows two fusions (overlap with the green and red signals). Single green and/or red signals indicate MYC rearrangements. Nuclei are counterstained with DAPI (blue). **(B)** Colony-formation assays were performed comparing PC3M-Cas9 cells to PC3M-Cas9 lines co-transfected with pairs of enriched sgRNA guides: (A169/A361) and (A169/ A423) following 14 days of treatment with 6μM MYCi975. Data was normalized to DMSO control and represented the mean ± S.D. from biological replicates. (*p<0.05; **p<0.01; ***p<0.001). **(C)** Relative cell viability of PC3M-Cas9 cells co-transfected with sgRNA pairs (A11/A362, A169/A423, and A11/A423) was measured following continuous exposure to MYCi975 at 6µM for 20 days, followed by 10µM until day 35. **(D, E)** Multiple-sequence alignment of the MYC eMBII and bHLH-LZ regions across vertebrate species. Blue dots denote DNA-binding residues in bHLH; green dots mark leucine-zipper dimerization residues. The MYC mutations identified are indicated in red font. **(F)** Expression and BLI Kd of recombinant MYC mutants WAL-QRA, WAL (eMBII mutant), and QRA (bHLH mutant) with MYCi975. **(G)** Colony formation assay of 22Rv1 and PC3 cells stably expressing Flag-tagged WT MYC and WAL-QRA MYC after treatment with DMSO and varying concentrations of MYCi975 for 14 days. Representative wells of colony formation assay are shown. **(H)** Cell viability of HO15.19 MYC and WAL-QRA MYC mutant was measured by IncuCyte live-cell imaging for 4 days with varying concentrations of MYCi975. **(I)** Immunoblot analyses of FLAG-tagged MYC protein stability in HO15.19 cells expressing WAL and QRA-MYC mutant, treated with 6µM MYCi975 for 24 hours. Actin serves as a loading control. **(J)** Quantification of FLAG-tagged MYC protein stability in HO15.19 cells expressing WT, WAL and QRA mutant, treated with 6µM MYCi975 for 24 hours. Data was normalized to DMSO control and represent the mean ± S.D. from biological replicates (n=3). (*p<0.05; **p<0.01; ***p<0.001).

**Figure S3.**
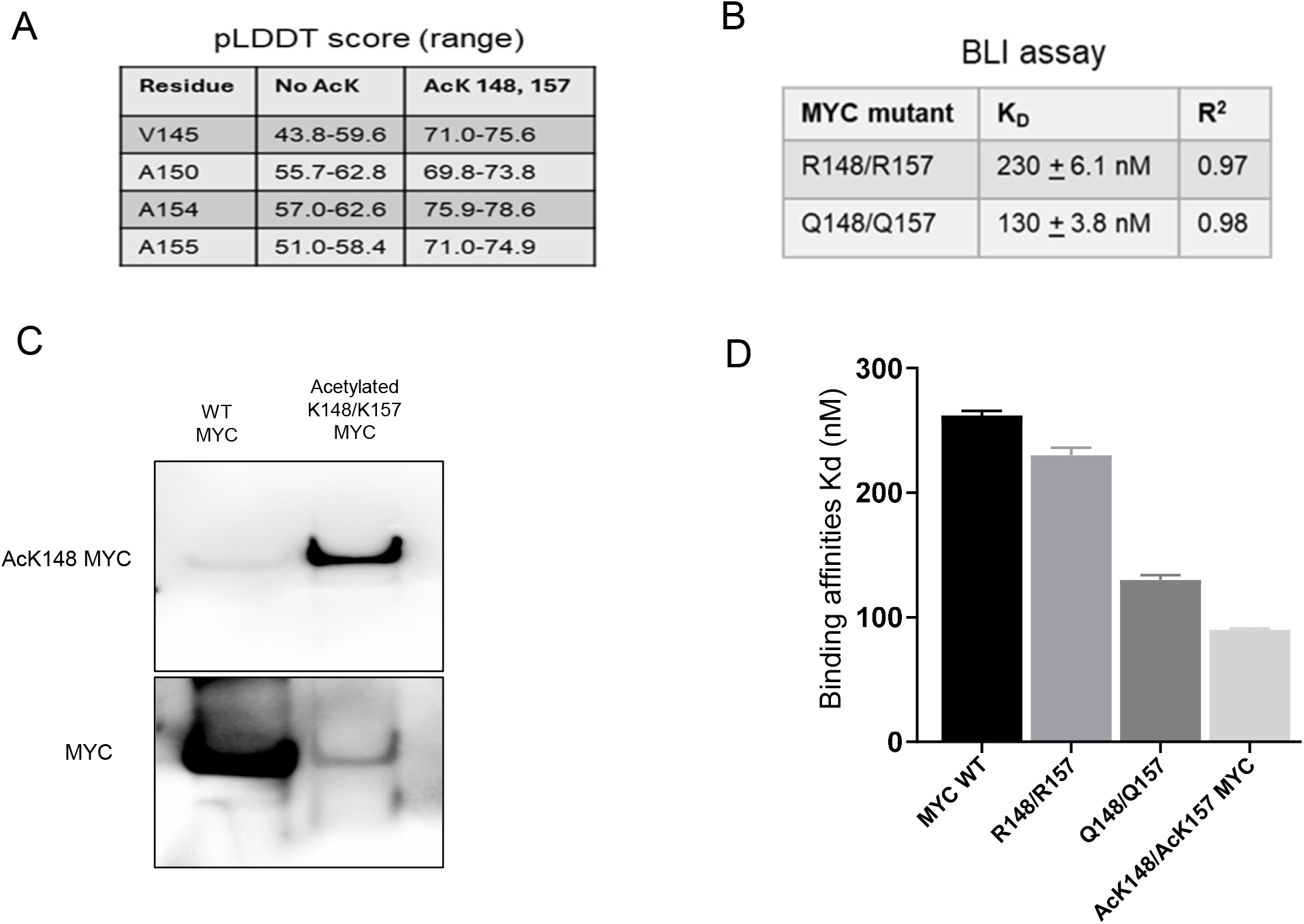
Acetylation of MYC lysine-148 and lysine-157 stabilizes the eMBII region and enhances MYCi975 binding. **(A)** Comparison of local pLDDT confidence scores from AlphaFold 3 structure predictions for the MYC eMBII region (residues V145–A155) in the absence (No Acetylation) and presence of acetylation at K148 and K157 (AcK148,157). Acetylation elevates scores into the high confidence range (>70) across all residues, indicating enhanced regional folding propensity. **(B)** Acetylation-mimic mutations increase MYCi975 binding affinity. BLI K_d_ of MYCi975 to deacetyl-mimic (R148/R157) and acetyl-mimic (Q148/Q157) MYC protein variants. **(C)** Validation of site-specific acetylation. Immunoblot analysis of purified recombinant MYC protein using an acetylation-specific antibody for K148 (AcK148) and a MYC antibody (Y69), confirming successful and specific acetylation at the K148 residue. **(D)** Comparison of BLI K_d_ for MYCi975 binding to recombinant WT, K148R/K157R, K148Q/K157Q, and AcK148/AcK157 MYC.

**Figure S4.**
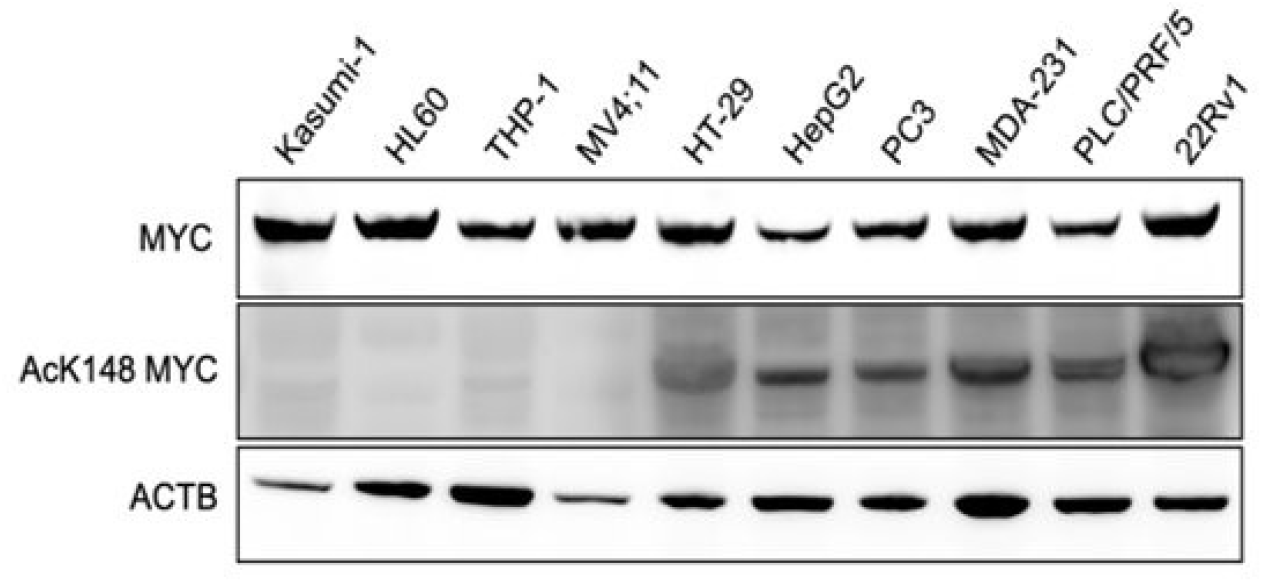
Immunoblot analyses showing MYC and AcK148-MYC protein expression across a panel of 10 cancer cell lines.

**Figure S5.**
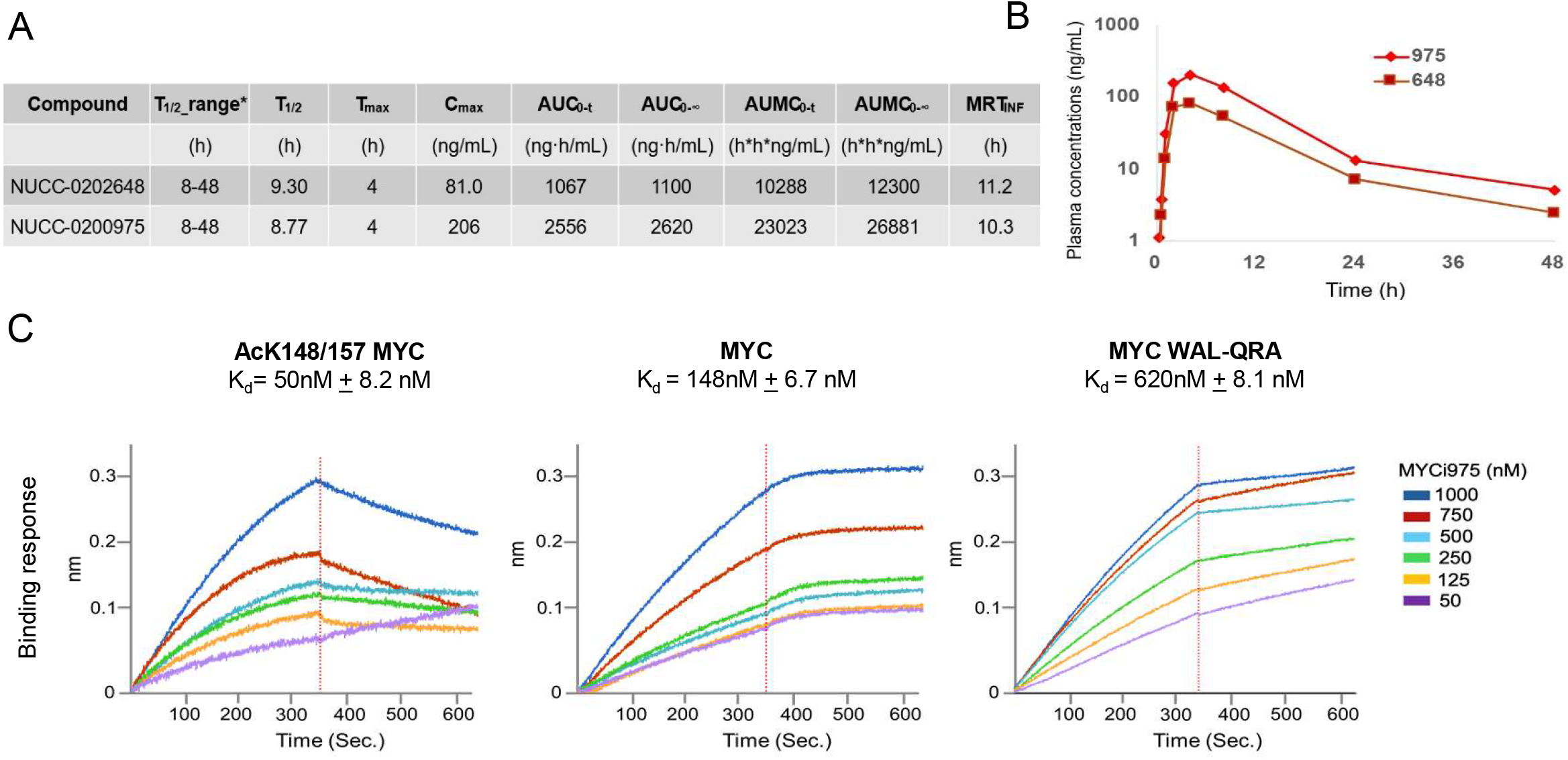
Pharmacokinetic properties and extended in vivo efficacy of MYCi648 compared with MYCi975. **(A)** Pharmacokinetic parameters of MYCi648 and MYCi975 in mice, including half-life (T_1/2_maximum concentration (C_max_), area under the curve (AUC), area under the first moment curve (AUMC), and mean residence time (MRT). **(B)** Plasma concentration-time profiles of MYCi648 and MYCi975. **(C)** BLI K_d_ of MYCi648 to acetylated MYC (AcK148/157-MYC), WT-MYC, and the WAL-QRA MYC mutant.

## Notes

### Competing Interest Statement

S.A.A. and G.E.S. are co-founders of Vortex Therapeutics, which is licensing of Intellectual Property (IP) related to the science and materials found in this manuscript. S.A.A., G.E.S, H.H., and M.I.T. are co-inventors on that same IP. All other authors declare that they have no competing interests.

### Summary of Updates

Nuclear magnetic resonance spectroscopy and molecular docking models of bHLH-LZ, eMBII and MYCi975 are added.

